# Morphological variation and strain identification of insects using wings and I^3^S

**DOI:** 10.1101/309468

**Authors:** Nayna Vyas-Patel, John D Mumford

## Abstract

Novel insect identification techniques often lead to speculation on whether the method could cope with any intraspecific variation that might occur in a species. Using I^3^S Classic (Interactive Individual Identification System, Classic) and images of mosquito wings, different mosquito strains were tested with a copy of the strain present or absent from the database which contained images of other strains of the test species. When a wing image of the exact species, strain and sex was present in the database, there was 100% (or near 100%) retrieval of the correct species and strain at rank one. When the exact strain was absent from the database, but other strains of the same species were present, the retrieval rates at rank one were again high (100%) in the majority of cases and when they were not, the correct species was generally retrieved at rank two. Out of 40 different species and strains tested, only three were significantly different at rank one when the exact strain was absent from the database. In general, images of field strains selected for each other and therefore were similar to each other in greater numbers and instances than for the laboratory strains tested. When a copy of a strain was absent from the database, but other strains/sibling species were present, I^3^S retrieved the correct strains/sibling species at rank one in the majority of cases. In the one case of transgenic mosquitoes tested, I^3^S could reliably be used to identify transgenic mosquitoes from the parent stock as they were retrieved 100% at rank one when both the transgenic and unmodified parent strains were present in the database. This indicates the potential of using I^3^S to distinguish transgenic or other selectively bred strains from a parent strain, also selectively bred and wild mosquitoes, at least in the first phase after field release. Similarly, hybrid strains, sibling species and members of species complexes as in the *Anopheles gambiae* species complex could also be correctly identified when copies of all the relevant species/strains/siblings were in the database. This contradicts the belief that only molecular characterisation could separate *A. gambiae* s.s. from *A. coluzzii*, or *A. arabiensis*; I^3^S could accurately separate them all. I^3^S worked as it was set up to do, retrieving closely resembling images of the test insects from the database and ranking them in order of similarity. Dealing with any intraspecific variation was therefore not an issue if the software (I^3^S) was used systematically. I^3^S complements molecular and traditional taxonomic methods for species identification and the separation of sibling complexes and strains. In future, it should become the norm to maintain databases of insect wings and other body part images for use in image recognition.

## Introduction

Morphological variation, large or small, exists in many organisms including insects and could potentially give rise to challenges in accurate species identification (Keeley 1982; Nosil & Reimchen, 2005; Ampuero et al 2009 & 2010 and Paz Garcia et al, 2015). This variation in appearance and size can occur over distances when a given species might present with differences in phenotype and/or size and is a feature of many island species that look different from their siblings on the mainland; or in the same species that live many miles or continents apart. It also occurs in species that experience different environmental conditions to their siblings living elsewhere. Termed ‘ecological variation’ it has been documented in many organisms including insects (Yi Bai et al 2016, Suman et al 2009, Dellicour et al 2017, Ekgachai et al 2013).

The wings of insects have long been known to be a reliable diagnostic feature for a given species (Woodward 1926, Wooton 1992, Wilke et al 2016). The use of algorithms and software to capture wing shape and the pattern and proportions of the vein structure on insect wings to differentiate species, means that even the smallest change in this shape, pattern and/or proportion can be detected quickly and easily. Image recognition software such as I^3^S Classic (the open source, Interactive Individual Identification System, I^3^S) can detect those changes far better than the human eye. Even if a test specimen was smaller or larger than the correct species present in the database, I^3^S takes account of the pattern and geometry of the veins on the test specimen to retrieve the closest reference image of a species, if not at rank one then at least within the first five ranks (Vyas-Patel et al 2016; Vyas-Patel & Mumford, 2017). Despite this use of mathematics, geometry and software, one of the criticisms for the use of technology and other ‘automated identification systems’ in insect species identification, has been the unverified argument (for I^3^S at least) that different strains of a given species might vary (intraspecific variation) leading to inaccuracies in species identification. Jean-Pierre Dujardin explained at great length why modern morphometric methods including image recognition, are capable of detecting even the smallest of change in phenotypic features that can exist between morphologically similar sibling species and aid the accurate identification of species and strain recognition (Dujardin, 2011).

The concept ‘different strains of a species’ is understood well when referring to the genotype where a ‘strain’ is a genetic variant or subtype of an organism. Different ‘strains’, ‘sibling species’ and ‘sub species’ are generally considered to be genetic variants of a given species. Variation can also occur in the phenotype and it was this ‘appearance based’, intraspecific variation within species that was of interest in this study. This can be assessed using geometric morphometric techniques or its modern equivalent - image recognition software. Insects respond to stress and conversely the lack of stress (very favourable conditions) in many different ways for example by changing their behaviour or by changes in the pattern of their life histories. The response may also result in changes in their morphology and physiology. It is increasingly being realized that it is not only distances or geographical isolation that can lead to phenotypic variation, but also factors such as nutrition (Pieterse et al 2017) and soil type (Benitez et al 2013). Pieterse et al (2017) noted that fruit flies reared on different fruits, notably apple and pear, had significantly different wing shapes compared to fruit flies reared on other fruits and that this could be detected and measured using geometric morphometric techniques to gauge wing shapes. Benitez (2013) demonstrated that using geometric morphometrics, the shape of the hind wings of *Diabrotica virgifera virgifera* changed according to the major soil types of Croatia. The wing morphology of two sibling *Drosophila* species was influenced by the different host plants (cactus) they fed on (Carreira et al 2006; Soto et al 2008 & 2010). Gomez-Cendra et al (2016) found that it was not geographical distance that affected the wing morphology of their fruit flies, but the host fruit available to the flies, those feeding on the same fruit had similar wing morphology no matter how far apart they lived. Using molecular markers and noting phenotypic differentiation, Gomez-Cendra et al (2016) reported that flies collected from and feeding on peaches or walnuts differed genetically and in appearance from each other regardless of geographical or temporal overlap. Yi Bai found that the grasshopper species *Trilophidia annulata* followed the Bergmann rule where individuals from higher, cooler latitudes with longer growing seasons were larger in size (using wing length as a measure) than individuals from the lower, warmer latitudes (Yi Bai et al 2016). There could be many other unknown factors that could potentially have an effect on phenotypic variation. Therefore, the present study considered any species collected from different locations as a ‘strain’, this included species reared in different laboratories and separate colonies within laboratories; if the same species was reared in different laboratories, they were considered as different strains of the species for the purposes of this study. Some of the species reared in laboratories were hybrids, created to prevent inbreeding, others were reared because they were known to be strains which were either resistant or susceptible to insecticides and/or susceptible or refractory to medically important parasites or viruses such as the malaria parasite or the Chikungunya, Zika and Dengue viruses. Both field caught and laboratory reared species and strains were examined. Mosquitoes were used as the test insect.

As evidenced above, a large body of information exists on the use of geometric morphometric methods to distinguish between insect species using wings. He-Ping Yang’s (2015) account of different geometric morphometric methods to identify insects using wing images currently provides the most encompassing review. Wilke et al (2016) dealt entirely with methods using wing morphometrics for the identification of different mosquito species. This comprehensive study is amongst the few to describe the use of software and image recognition for the identification of species and strains. The project used the freely downloadable software, I^3^S Classic to test the identification of different strains of insect species (both laboratory reared and field caught) to determine if it could retrieve the correct species and strain when a reference image of the test taxon was not in the database but other strains of the test species were.

Ordinarily, as I^3^S is very accurate at retrieving an image of the test species if it is present in the database (Vyas-Patel & Mumford 2017), there would be no point in trying to retrieve a copy of an image which was not present in the database. In such cases, only the closest match could be retrieved and the score attached to each ranked image in the results would give some indication as to how far the match at rank 1 was from the test. Knowledge of the contents of the database was emphasised in previous studies using I^3^S, as the software cannot fully match an image of a species which was not present in the database (Vyas-Patel & Mumford 2017); only the closest match could be retrieved. However, for the purpose of this study, example images of all the species and strains to be tested were first uploaded into the database (one wing image of each species, strain and sex) and each species and strain was tested. Next, the test strain reference was removed from the database, leaving other, different strain/s of the same species in the database and re-tested. Hence strains were tested with and then without a copy of the specific reference strain in the database.

Most image recognition software is designed to compare like for like (similarities) and rank the results. Hence, even if a specific reference image was present in the database, if the test had not been aligned in the same way as the database image of the same species, or marked in the same way, it could result in errors. Care was taken to align and mark the wings consistently, in the same way for both the test and database images. Alignment was achieved by eye using software to rotate the image by degrees (custom rotate, Photoshop), pin point precision was not required. The wing samples used were largely intact (unbroken) except in the few cases of field caught specimens where the numbers available for testing were low. In such cases, imperfect, broken wings were also imaged and tested. Any major folds if present were gently eased out by the placement of a coverslip over the wing before a photograph was taken. This was particularly important for smaller wings such as in mosquitoes, which are generally in the order of 3mm long, but less so for larger insect wings, where the stronger vein structure tended to keep the wings flat (experience from previous studies using Hymenopteran wings).

## Materials and Method

### Insect Specimens

Mosquito strains originating from around the world and reared in different laboratories, as well as field caught strains from different locations were used for the study. The species were identified by trained scientists in every case. The donors and the available metadata for each strain are given in Table 1 together with footnotes.

**Table 1.** Strains and species of Mosquitoes used with available metadata.

|  |  |
| --- | --- |
| *Aedes aegypti* IAEA | Originating from Brazil, laboratory reared at the International Atomic Energy Agency (IAEA), Seibersdorf, Austria. |
| *Aedes aegypti* | From West Africa, insecticide susceptible, refractory to |
| LSHTM | Filarial infections. Laboratory reared since 1926, London. |
| *Aedes aegypti* UG | Field caught, University of Georgia, USA. |
| *Aedes albopictus* UG | Field caught from Fayetteville, Alabama, by University of Georgia, USA. |
| *Aedes albopictus* SAV | Field caught from Savannah, Georgia, USA. |
| *Aedes albopictus* IAEA | Originating from Rimini, Italy and laboratory reared at IAEA. |
| *Aedes vexans* UG | Field caught from Clinton, South Carolina, Laurens County, Ex River Bottom, by University of Georgia, USA. |
| *Aedes vexans* Den | Field caught from Denver, USA, +40.18920N, -105.05975W, Alt. 4,995 ft. |
| *Aedes vexans* | Field caught from the Atlanta, Georgia region, USA. |
| *Anopheles arabiensis* IAEA | Originating from Dongola, Sudan, Africa. Laboratory reared at IAEA and used in the Sterile Insect Technique, SIT. |
| *Anopheles arabiensis* ICL | Obtained from MR4, BEI resources (Malaria Research & Reference Reagent Resource Centre), the strain was laboratory reared at Imperial College London (ICL) and originates from Dongola, Sudan. |
| *Anopheles gambiae* G3, ICL | Laboratory reared at Imperial College London (ICL), UK, the strain G3 ICL, was obtained from MR4, BEI resources. Originally, the strain was collected from McCarthy Island, The Gambia, West Africa, but is now known to be a hybrid. |
| *Anopheles gambiae* G3, LSHTM | Laboratory reared at the London School of Hygiene & Tropical Medicine (LSHTM) since 1998 for its susceptibility to Dieldrin. |
| *Transgenic Cas9* (from *Anopheles gambiae* G3) | The Cas9 transgenic mosquitoes were genetically modified using the G3 strain at Imperial College London (ICL) and selected for fluorescent marker expression at each generation, (Hammond et al 2016). |
| *Anopheles gambiae* | Laboratory reared at LSHTM from 1998 onwards, from |
| *Kisumu* | Kisumu, Kenya, Africa. Fully susceptible to Pyrethroids. |
| *Anopheles gambiae* Tia (Tiassale) | Laboratory reared at LSHTM from 2013 onwards. From Tiassale, Cote d'Ivoire, Africa. |
| *Anopheles gambiae* N'guesso (*A. coluzzii*) ICL | Colonised in 2006 at ICL, from mosquitoes collected around Yaoundé Cameroon and reclassified as *A. coluzzii* following molecular characterisation, Habtewold et al, 2016. |
| *Anopheles gambiae* N'guesso (*A. coluzzii*) LSHTM | Laboratory reared at LSHTM since 2013, this strain was obtained from ICL (not known at the time of testing). |
| *Anopheles gambiae* Keele (*A. coluzzii*) | Laboratory reared at the University of Glasgow. Originally a hybrid colony from crosses between four different *A. gambiae* colonies maintained at Keele University; molecular characterisation indicated that the Keele colony was *A. coluzzii* (Ranford-Cartwright, 2016). |
| *Anopheles stephensi* SDA500 | Laboratory reared at LSHTM from a mixture of stocks from Sind province, Pakistan. |
| *Anopheles stephensi* Red | Maintained at LSHTM, from a colony with genetically recessive red eyes. |
| *Anopheles stephensi* Beech | From New Delhi India, colonised since 1947 at Beecham's laboratory and subsequently reared at the LSHTM. |
| *Anopheles stephensi* STML | Laboratory reared at LSHTM. From Lahore, Pakistan, colonised since 1979. |
| *Anopheles stephensi* Dub5 | Laboratory reared at LSHTM. From Dubai, colonised since 1989. |
| *Coquillettidia perturbans* UG | Field collected by University of Georgia (UG) entomologists from the Georgia area, USA. |
| *Coquillettidia perturbans* Sav | Field collected by the Centre for Disease Control (CDC) entomologists, from Savannah, Georgia, USA. |
| *Coquillettidia perturbans* Den | Field collected from Denver, +39.93760N,-105.05740W, Alt 5,380 ft. |
| *Coquillettidia* | Field collected from the Atlanta region (Atl), Georgia, |
| *perturbans* Atl | USA. |
| *Culex quinquefasciatus* Mu | Laboratory reared at the LSHTM, pyrethroid resistant.<br>From Muheza (Mu) Tanzania. |
| *Culex quinquefasciatus* TPRI | From a colony maintained at the Tropical Pesticide Research Institute (TPRI), Tanzania. Reared at LSHTM since 2010. |
| *Culex quinquefasciatus* Sav | Field collected from the Savannah (Sav) region, USA. |
| *Culex quinquefasciatus* UG | Field collected from Clarke County by University of Georgia (UG) Athens, Georgia, USA. |
| *Ochlerotatus taeniorinchus* Sav | Field collected from the Savannah, Georgia region, USA. |
| *Ochlerotatus taeniorinchus* UG | Field collected from Darien, McIntosh County, Georgia, by University of Georgia entomologists, USA. |
| *Ochlerotatus triseriatus* Sav | Field collected from the Savannah, Georgia region, USA. |
| *Ochlerotatus triseriatus* UG | Field collected from Clarke County, Georgia, by University of Georgia entomologists, USA. |
| *Ochlerotatus sollicitans* Sav | Field collected from the Savannah, Georgia region, USA. |
| *Ochlerotatus sollicitans* UG | Field collected from Darien, McIntosh County, Georgia, by University of Georgia entomologists, USA. |
| *Psorophora ferox* Sav | Field collected from the Savannah, Georgia region, USA. |
| *Psorophora ferox* UG | Field collected by University of Georgia, Athens, Georgia, USA. |

### Preparation and marking of the Images

The wings were dissected from the body under a standard dissection microscope, photographed with a Samsung NV10 digital camera, using only the sub-stage lighting of the microscope, as this produced a clear image of the wing shape and venation. Prior to taking a photograph, a clean microscope cover slip was placed on the wing to ensure that any folds were gently flattened out. Each image was uploaded into an Adobe Photoshop (CS6) image editor and rotated so that the point of insertion of the wing into the body of the insect always faced to the left and the wing was aligned to be as horizontal as possible, using ‘Image rotate’ and ‘Custom rotate’ in the top menu bar of Photoshop. The newly aligned and rotated images were saved as .jpg files, creating a different file for each species and sex.

I^3^S Classic, version 4.02, was downloaded from the internet and a ‘fingerprint’ (.fgp) image of each wing image was prepared and stored in the database as described in previous studies (Vyas-Patel & Mumford 2017) and shown in Figure 1. Any clearly visible point and landmark wing feature may be chosen for marking as long as subsequent images were also consistently marked in the same way. Each fingerprint file was saved and could then be used either for reference database creation or as a test image. A comprehensive guide to the use of I^3^S and how images can be prepared, stored and analysed is given in the instructions on the I^3^S website together with a tutorial (http://www.reijns.com/i3s/download/I3S%20Classic.pdf).

**Figure 1:**
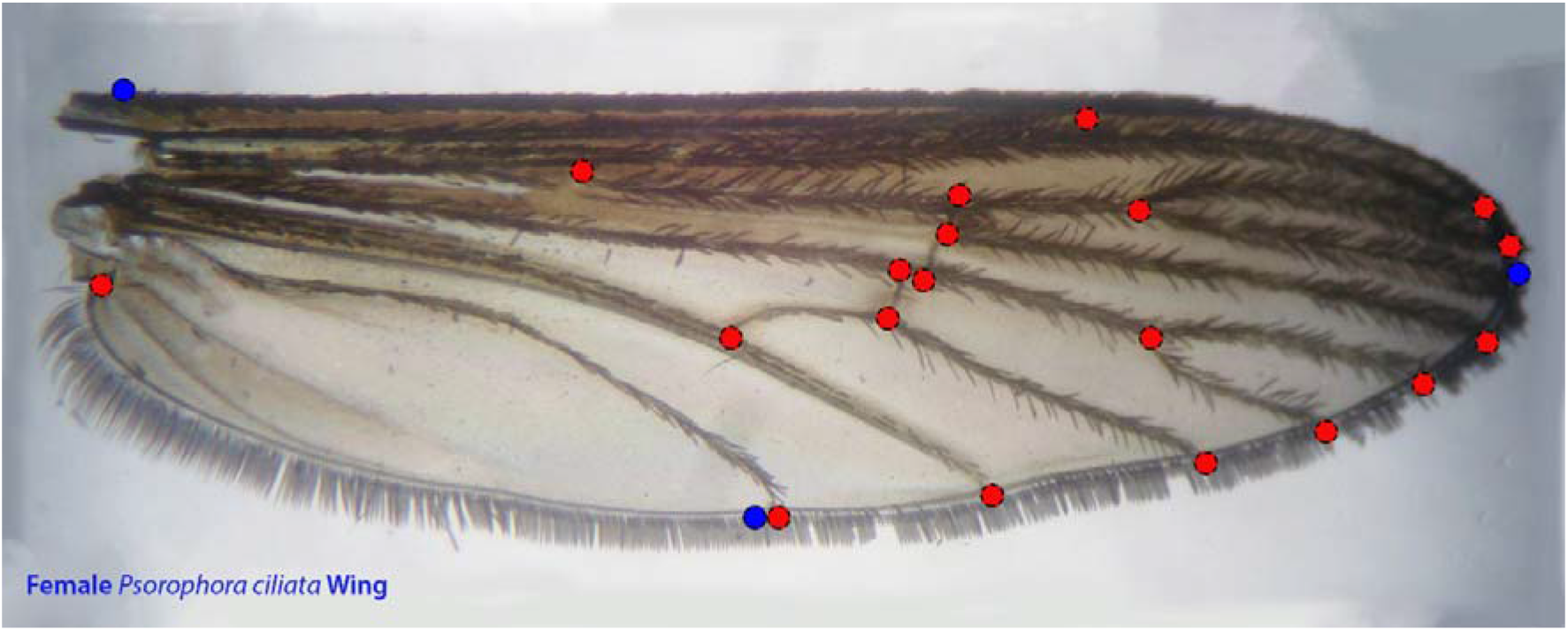
A mosquito wing indicating the markings made using I^3^S.

### Database creation

A database of fingerprint (.fgp) files was created. This comprised one image of each sex and strain of *Aedes aegypti* (both field caught and laboratory reared), *Aedes albopictus* (field caught and laboratory reared), *Aedes vexans* (field caught), *Anopheles arabiensis* (laboratory reared), Transgenic Cas9 derived from hybrid *Anopheles gambiae* G3 reared at Imperial College London (ICL), *Anopheles gambiae* G3 (ICL) *Anopheles gambiae* (Kisumu & Tiassale strains), *Anopheles gambiae N’guesso (A. coluzzii), Anopheles gambiae Keele (A. coluzzii), Anopheles stephensi* (laboratory reared, strains Red, Beech, SDA500, STML and Dub5), *Coquillettidia perturbans* (field caught), *Culex quinquefasciatus* (field caught and laboratory reared - Muheza, TPRI, UG), *Ochlerotatus taeniorinchus* (field caught), *Ochlerotatus triseriatus* (field caught), *Ochlerotatus sollicitans* (field caught), and *Psorophora ferox* (field caught). A total of forty different strains and sixty different wing images of mosquito strains were uploaded into the database. Wing images of both sexes were included from the laboratory reared species/strains. In the case of field caught specimens only females were available, as the traps were designed to attract adult female mosquitoes and so largely collected females. Furthermore, in the case of field caught specimens where the numbers available were low, broken edged (imperfect) wings were also used. Images were added and removed from the database as required for each of the comparisons. The test images were therefore assessed with and without a corresponding reference wing image in the database and rank 1 and 2 results were noted in each case, Table 2.

**Table 2.** Total percentage of strains correctly identified at Ranks 1 & 2, with and without an image of the test strain in the database (DB).

| Test Strain | R1 Test in DB | R2 Test in DB | R1 Test **not** in DB | R2 Test **not** in DB |
| --- | --- | --- | --- | --- |
| *Aedes aegypti* -Aae IAEA (15M, 15F) | 100% Aae IAEA | 100% Aae Strains (53% UG, 47% LSHTM) | 96% Aae Strains (72% UG, 24% LSHTM), 4% Aalbo | 88% Aae Strains (62% LSHTM, 26% UG), 12% Aalbo |
| *Aedes aegypti* – Aae LSHTM (15M, 15F) | 100% Aae Strains (80% LSHTM, 16% UG, 4% IAEA) | 100% Aae Strains (84% UG, 16% IAEA) | 100% Aae Strains (73% UG, 27% IAEA) | 80% Aae Strains (54% IAEA, 26% UG), 7% Mu, 13% Coq p |
| *Aedes aegypti* – Aae UG (15F) | 100% Aae Strains, (80% UG, 20% LSHTM) | 100% Aae Strains (55% LSHTM, 20% Aae UG, 25% Aae IAEA) | 100% Aae (45% IAEA, 55% LSHTM) | 75% Aae Strains (38% LSHTM, 37% IAEA), 12% Cq Mu, 13% Coq p |
| *Aedes vexans* -Avex Atl (30F). | 97% Avex Strains (51% Atl 40% UG, 6% Den), 3% Aalbo | 85% Avex Strains (45% UG%, 40% Den), 15% Aae LSHTM | 85% A vex Strains (45% Den, 40% UG) 15% Aae LSHTM | 75% Avex Strains (35% Den, 40% UG), 25% Aae LSHTM |
| *Aedes vexans* – Avex Denver (30F) | 97% Avex Strains (44% Atl, 33% Den, 20% UG), 3% Aae LSHTM | 82% Avex Strains (34% UG, 20% Den, 28% Atl), 9% Aae LSHTM, 9% Otaen | 96% Avex Strains (53% Atl, 43% UG), 4% Aae LSHTM | 67% Avex Strains (38% Atl, 29% UG), 14% Otaen, 19% Aae |
| *Aedes vexans* – Avex UG | 100% Avex Strains (31% | 100% Avex Strains (62% | 83% Avex Strains (55% Atl, 28% | 61% Avex Strains (33%Atl, 28% Den), |
| (15F) | Den, 54% UG<br>15% Atl). | Atl, 23% Den,<br>15% UG) | Den) 14% Aae,<br>3% Otaen | 16% Aae, 23%<br>Otaen |
| *Aedes albopictus* -<br>Aalbo IAEA<br>(15M, 15F) | 92% Aalbo<br>Strains (77%<br>IAEA, 15%<br>Sav)<br>8% AaeUG | 75% Aalbo<br>Strains (67%<br>Sav, 8% IAEA),<br>17% Aae, 8%<br>Otriseri | 75% Aalbo<br>Strains (54% Sav,<br>21% UG), 10%<br>Aae 15% Otriseri | 39% Aalbo strains<br>(31% Sav, 8% UG),<br>38% Aae, 23%<br>Otriseri |
| *Aedes albopictus*<br>University of<br>Georgia –<br>Aalbo UG<br>(16F) | 100% Aalbo<br>Strains (100%<br>UG) | 81% Aalbo<br>Strains (81%<br>Sav) 19% Aae | 81% Aalbo strains<br>(62% IAEA, 19%<br>Sav), 19% Aae | 56% Aalbo Strains<br>(36% IAEA, 20%<br>Sav), 18% Avex,<br>26% Aae |
| *Aedes Albopictus*<br>Savannah -<br>Aalbo Sav’<br>(10F) | 100% Aalbo<br>Strains (80%<br>Sav 20%<br>IAEA) | 80% Aalbo<br>Strains (60%<br>UG, 20% Sav),<br>20% Aae | 80% Aalbo<br>Strains (50%<br>IAEA, 30% UG),<br>Avex 20% | 40% Aalbo strains<br>(25% AEA, 15%<br>UG), 38% Aae, 22%<br>Avex |
| *Anopheles arabiensis* -<br>Aarab, IAEA<br>(8M, 8F) | 100% Aarab<br>Strains (100%<br>IAEA) | 77% Aarab<br>Strains (77%<br>ICL), 15% Agam<br>G3, 8% Agam<br>Tia<br>**(100% Agam**<br>**Complex)** | 76% Aarab<br>Strains<br>(76% ICL) 12%<br>Agam Tia, 12%<br>Agam G3 **(100%**<br>**Agam Complex)** | 40% Agam G3, 20%<br>Agam Tia, 20%<br>Keele, 20% Keele<br>**(100% Agam**<br>**Complex)** |
| *Anopheles arabiensis* –<br>Aarab ICL<br>(7M, 7F) | 100% Aarab<br>Strains (75%<br>ICL, 25%<br>IAEA) | 75% Aarab<br>IAEA, 17%<br>Agam G3, 8%<br>Ngu **(100%**<br>**Agam**<br>**Complex)** | 75% Aarab<br>(IAEA) 17%<br>Agam G3, 8%<br>Ngu **(100% Agam**<br>**Complex)** | 84% Agam Tia, 8%<br>Agam G3, 8% As<br>STML<br>**(92% Agam**<br>**Complex)** |
| *Anopheles gambiae* – Agam G3 LSHTM (15M, 15F) | 100% Agam G3 LSHTM Strains | 39% Agam Kisumu, 29% Keele, 4% Agam Tia, 26% Agam Ngu, 2% As Red **(98% Agam complex Strains)** | 14% Agam Kisumu 36% Keele, 36% Aarab, 7% Tia, 7% As Dub5 **(93% Agam Complex)** | 27% Keele, 19% Agam Kisumu, 12% Agam Ngu, 12% Agam Tia, 8% Aarab, 3% As Dub5, 10% As Red, 9% As Beech **(78% Agam Complex)** |
| *Anopheles gambiae* – Agam G3 ICL (10F, 10M) | 71% Agam G3 ICL Strain, 29% Keele **(100% Agam complex)** | 7% Agam G3 ICL, 39% Keele, 39% Aarab, 8% Agam Tia, 7% As Dub5 **(93% Agam Complex)** | 72% Keele, 14% Aarab, 7% Agam Tia, 7% As Beech **(93% Agam complex Strains)** | 72% Aarab, 21% As Dub5, 7% As STML **(72% Agam Complex Strains)** |
| Transgenic Cas9 – Trans (gene edited Agam G3 ICL), (10M, 10F) | 100% Trans | 67% G3 ICL, 33% Aarab **(100% Agam Complex Strains)** | 67% G3 ICL, 33% Aarab **(100% Agam Complex Strains)** | 13% G3 ICL, 27%, Keele, 42% Aarab, 8% Dub5, 10% As STML **(82% Agam Complex Strains)** |
| *Anopheles gambiae* – Agam Kisumu (20M, 20F) | 100% Agam Kisumu | 25% Aarab, 35% Keele, 8% Ngu, 10% Tia, 20% G3 (LSHTM), 2% Dub5 **(88% Agam Complex Strains)** | 30% Aarab, 28% G3 (LSHTM), 30% Keele, 3% Ngu, 6% Tia, 3% Dub5 **(97% Agam Complex strains)** | 36% Aarab, 36% G3 (LSHTM), 9% Keele, 7% Tia, 12% Ngu **(100% Agam Complex strains)** |
| *Anopheles gambiae* – Agam Tia | 100% Agam Tia | 21% Aarab, 26%, Keele, 16% Kis, 33% | 27% G3, 29%, Aarab, 18% Kis, 26% Ngu **(100%** | 22% G3, 25% Aarab 20% Keele, 33% Kis **(100% Agam** |
| (Tiassale)<br>(15M, 15F) |  | Ngu, 4% G3<br>**(100% Agam Complex)** | **Agam Complex)** | **Complex)** |
| *Anopheles gambiae* N'guesso LSHTM (A coluzzii hybrid) (15M, 15F) | 100% Agam Ngu LSHTM | 76% Ngu ICL, 14% Keele, 3% G3 7% Dub5<br>**(93% Agam Complex)** | 60% Ngu ICL, 33% Keele, 3% As Red, 4% As Beech **(93% Agam Complex)** | 20% Ngu ICL, 14% Aarab, 21% Tia, 20% Keele, 18% As Red, 7% As Beech **(75% Agam Complex)** |
| *Anopheles gambiae* N'guesso ICL (A coluzzii hybrid) (10M, 10F) | 100% Agam Ngu ICL | 53% Ngu LSHTM, 39% Keele, 4% G3, 4% Tia<br>**(100% Agam Complex)** | 65% Ngu LSHTM, 35% Keele **(100% Agam Complex)** | 16% Ngu LSHTM, 22% Tia, 13% As Red, 13%, Aarab Imp, 13% Keele 13% As STMI, 10%, As Red **(77% Agam Complex)** |
| *Anopheles gambiae* Keele (A coluzzii hybrid) (15M, 15F) | 82% Keele, 9% Ngu, 9% Aarab **(100% Agam Complex)** | 18% Keele, 45% Ngu, 17% G3, 5% Dub5, 15% Aarab<br>**(95% Agam Complex)** | 50% Ngu, 23% Aarab, 27% G3 **(100% Agam Complex)** | 15% Ngu, 40% Aarab, 14% G3, 18% Tia, 8% Dub5, 5% As Red **(87% Agam Complex)** |
| *Anopheles stephensi* - As SDA500 LSHTM (16M, 16F) | 82% As SDA500 6% As Red, 6% As Beech, 6% As STML, **100% As strains** | 40% As Beech, 12% As Dub5, 36% As Red, 12% As STML, **100% As Strains** | 38% As Red, 26% As Beech, 36% As Dub5, **100% As Strains** | 40% As Beech, 12% As Dub5, 36% As Red, 12% As STML, **100% As Strains** |
| *Anopheles stephensi* - As Red LSHTM | 100% As Red | 34% As SDA500, 44% As Beech, 22% | 35% As SDA500, 42% As Beech, 15% As Dub5, | 44% As Beech, 34% As SDA500, 22% Agam Ngu, **78% As** |
| (15M, 15F) |  | As Dub5 **100% As Strains** | 8%Ngu **92% As Strains** | **Strains** |
| *Anopheles stephensi* - As Beech LSHTM (15M, 15F) | 60% As Beech 30% As Red, 10% As SDA500 **100% As strains** | 31% As Beech, 22% As SDA500, 39% As Red, 8% Aarab. **92% As Strains** | 23% As SDA500, 61% As Red, 8% As STML, 8% Aarab. **92% As Strains** | 45% As SDA500, 31% As Red, 8% As Dub5, 8% Aarab, 8% Agam Ngu. **84% As Strains** |
| *Anopheles stephensi* – As STML LSHTM (15M, 15F) | 87% As STML, 13% As Red. **100% As Strains** | 3% As STML, 40% As Red, 35% As SDA500, 22% As Beech. **100% As Strains** | 40% As SD500, 22% As Beech, 23% As Red, 3% Agam Kis, 4% G3, 8% Tia. **85% As Strains** | 44% As SDA500, 25% As Beech, 15% Agam Kis, 16% Agam G3. **69% As Strains.** |
| *Anopheles stephensi* – As Dub5 (15M, 15F) | 67% As Dub5, 33% As Red. **(100% As Strains)** | 42% As Red, 35% As Beech, 16% As SDA500, 7% As STML **(100% As Strains)** | 42% As Red, 35% As Beech, 16% As SDA500, 7% As STML **(100% As Strains)** | 25% As Beech, 18% SDA500, 30% As Red, 18% Agam Kis, 9% Agam Ngu **(73% As Strains)** |
| *Coquillettidia perturbans* Coq p UG (16F) | 100% Coq p. (32% UG, 56% Den, 12% Atl) | 100% Coq p (31% Atl, 31% Den, 32% UG, 6%, Sav) | 94% Coq p (56% Atl 38% Sav), 6% Avex | 92% Coq p (44% Atl, 48% Sav), 8% Avex |
| *Coquillettidia perturbans* Coq p Sav (14F) | 100% Coq p (Sav 59%, 41% UG) | 100% Coq p (25% Sav, 25% Den, 25% Atl, 25% UG) | 100% Coq p (42% Atl, 42% Den 16% UG) | 100% Coq p (42% Atl, 42% Den, 16% UG) |
| *Coquillettidia perturbans* Coq p Den | 100% Coq p (Den) | 100% Coq p (72% Atl 14%, Sav 14%) | 100% Coq p (86% UG, 14% Sav) | 100% Coq p (72% Atl 14% Sav, 14% UG) |
| (15F) |  | UG) |  |  |
| *Coquillettidia perturbans* Coq p Atl (15F) | 100% Coq p (72% Atl, 14% UG, 14% Den) | 100% Coq p (6% Atl, 35% UG, 45% Den, 14% Sav) | 100% Coq p (43% Den, 21% Sav, 36% UG) | 100% Co q (29% Sav, 42% Den, 29% UG) |
| *Culex quinquefasciatus* Cq Muheza LSHTM (15M, 15F) | 100% Cq Strains (71% Mu, 21% TPRI, 8% Sav) | 71% Cq Strains (29% Mu, 42% TPRI), 29% Coq p Atl | 71% Cq Strains (44% TPRI, 27% Sav), 15% Coq p, 14% Avex | 33% Cq Strains (16% TPRI 17% Sav), 13% Cq tar, 13% Aae, 25% Avex 8% Coq p 8% |
| *Culex quinquefasciatus* TPRI (10M, 10F) | 100% Cq Strains (80% TPRI 20% Mu) | 100% Cq Strains (20% TPRI, 58% Mu, 22% Sav) | 84% Cq Strains (38% Mu, 46% Sav), 11% Avex, 5% Aae | 48% Cq Strains (20% Mu, 28% Sav) 18% Coq p, 20% Aae, 14% Avex |
| *Culex quinquefasciatus* Savannah (5F) | 100% Cq Strains (Sav 67% Mu 22% TPRI 11%) | 66% Cq Strains (44% TPRI, 22% UG) 22% Aae, 12% Avex | 66% Cq Strains (22% UG, 22% Mu, 22% TPRI) 24% Aae, 10% Avex | 55% Cq Strains (22% Ug, 22% Mu, 11% TPRI), 12% Avex, 33% Aae |
| *Culex quinquefasciatus* UG (5F) | 100% Cq strains (50% Sav, 25% Mu, 25% UG) | 75% Cq Strains (50% Sav, 25%, UG) 25% Avex | 85% Cq Strains (50% TPRI, 10% Mu 25% Sav), 15% Avex | 40% Cq Strains (20% TPRI, 20% UG), 30% Avex, 30% Aae |
| *Ochlerotatus taeniorhynchus* – Otaenor Savannah (6F) | 100% Otaenor Strains (80% Sav, 20% UG) | 70% Otaenior Strains (40% UG, 30% Sav) 20% Avex, 10% Aae | 60% Otaenior Strains (60% UG), 30% Avex, 10% Aae | 35% Otaenior strains (35% UG), 40% Avex, 25% Aae |
| *Ochlerotatus taeniorhynchus* UG (6F) | 100% Otaenior (UG 100%) | 55% Otaenior (55% Sav), 33% Otriseri 12% | 55% Otaenior (Sav 55%) 33% Otriseri 12% Avex | 25% Otaenior (25% Sav), 18% Otrisei, 57% Avex |

|  |  | Avex |  |  |
| --- | --- | --- | --- | --- |
| *Ochlerotatus triseriatus* Otriseri Savannah (6F) | 100% Otriseri Sav | 60% Otriseri (60% UG) 25% Otaenior, 15% Aae | 60% Otriseri (60% UG), 25% Otaenior 15%, Aae | 50% Avex, 50% Aae, 0% Otriseri |
| *Ochlerotatus triseriatus* UG (6F) | 100% Otriseri (UG 100%) | 43% Otriseri Strains (43% Sav), Avex 57% | 43% Otriseri Strains (43% Sav), 57% Avex | 72% Avex, 14% Otaenior, 14% Aae, 0% Otriseri |
| *Ochlerotatus sollicitans* Osoll UG (7F) | 100% Osoll UG | 77% Osoll Sav, 23% Avex | 77% Osoll Sav, 23% Avex | 29% Osoll Sav, 29%, Avex, 13% Otriseri 29% Otaenior |
| *Ochlerotatus sollicitans* Sav (7F) | 100% Osoll Sav | 100% Osoll UG | 100% Osoll UG | 60% Aae Ishtm, 40% Otaenior, 0% Osoll |
| *Psorofera ferox* – Pferox Savannah (6F) | 100% Pferox Strains (80% UG, 20% Sav) | 100% Pferox Strains (Sav 80%, UG 20% ) | 80% Pferox Strains (80% UG) 20% Aae Ishtm | 40% Otaenior, 60% Aae Ishtm, 0% Pferox |
| *Psorofera ferox* UG (6F) | 100% Pferox UG | 100% Pferox Sav | 100% Pferox Sav | 40% Avex Den, 40% Otriseri, 20% Otaenior, 0% Pferox |

M = Numbers of males used, F = Numbers of females used; DB= database.
R1 & R2 = Rank 1 and Rank 2 correctly identified strains
2. There was only one reference image of each strain in the database and once this had been selected at rank 1, it was not possible for it to be selected again at rank 2, hence the 0% correct strains at rank 2 when rank 1 had 100% correctly identified strain results (no further correct strains could be selected at rank 2).
3. In cases where strains were part of a species complex, the total percentage of sibling strains/complexes selected is highlighted in bold.

Any significant difference in results when the test image was present compared to absent from the database is given in Table 3. The salient features of the observed results for each strain is given in Table 4.

**Table 3:**
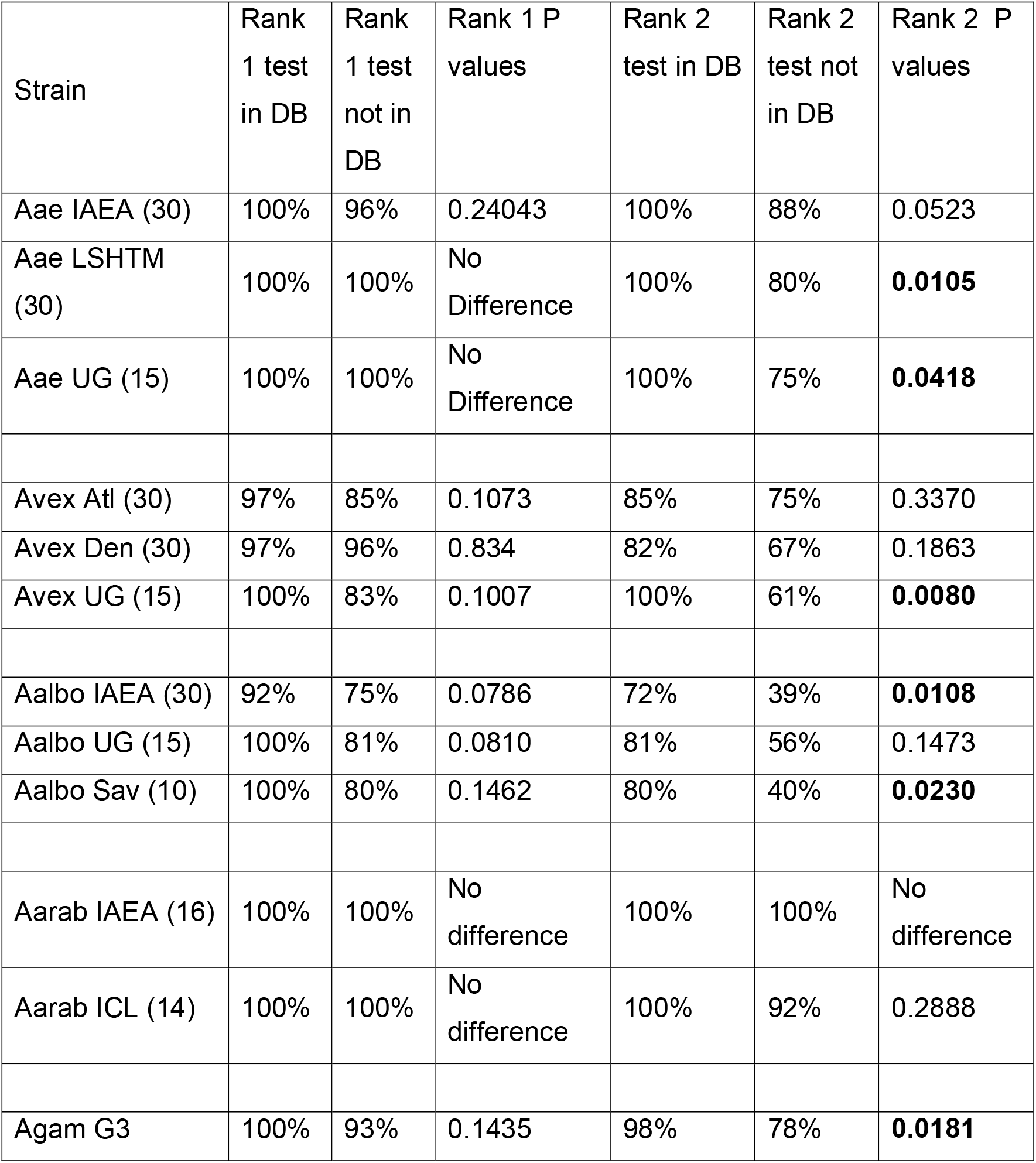
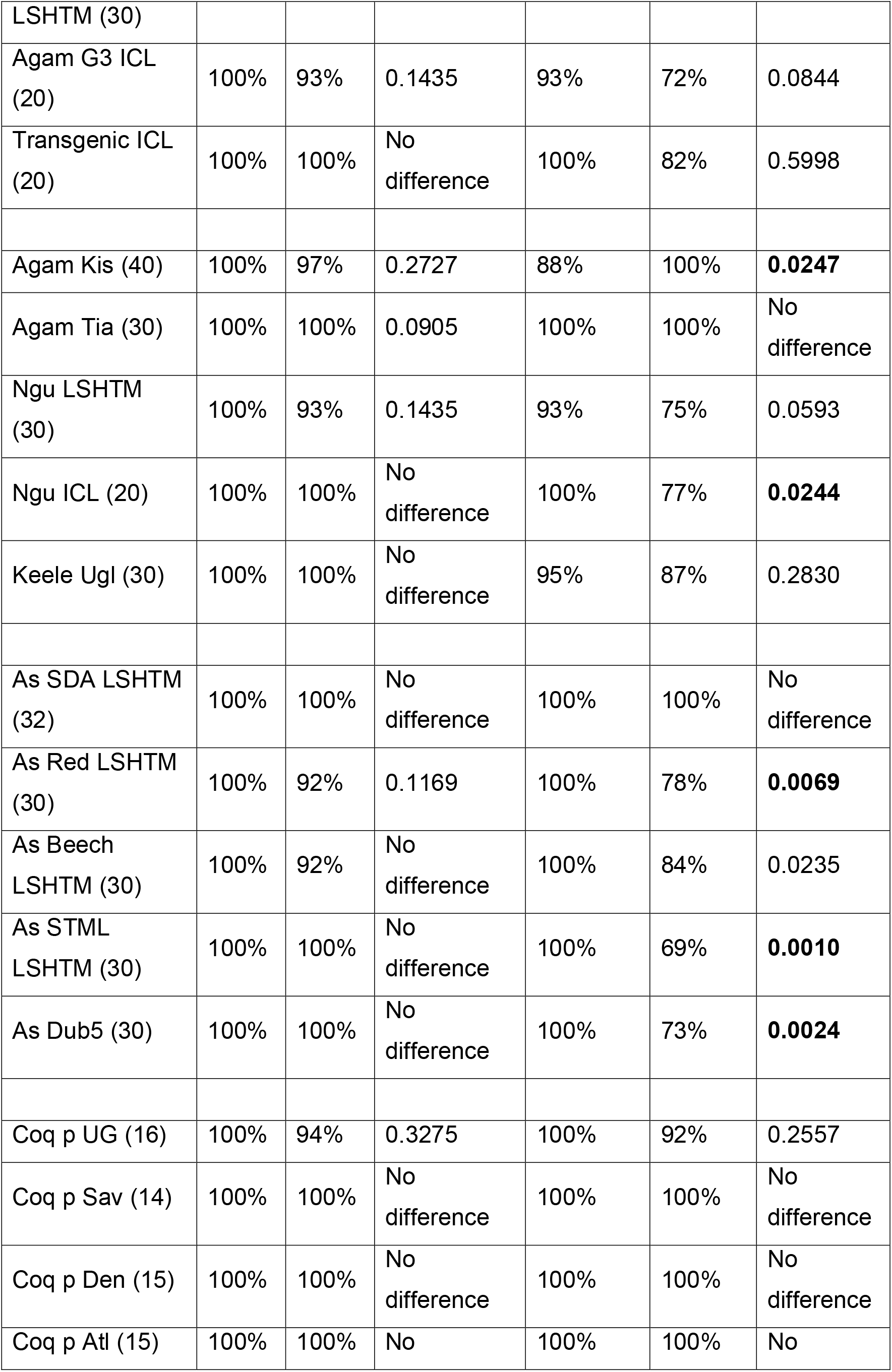
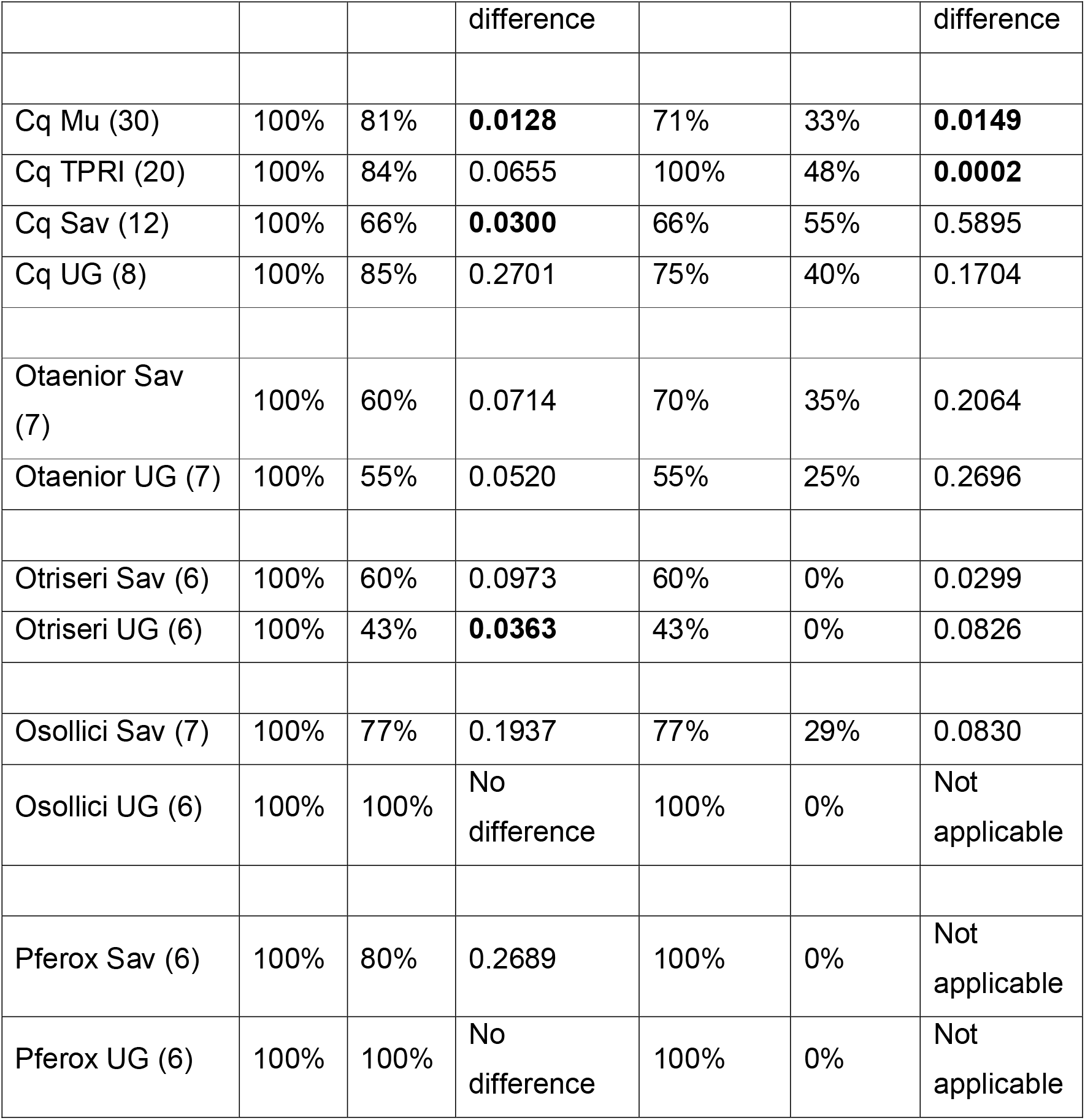
The difference between the percentages of correctly identified strains in the presence and absence of the test strain in the database.

| Strain | Rank 1 test in DB | Rank 1 test not in DB | Rank 1 P values | Rank 2 test in DB | Rank 2 test not in DB | Rank 2 P values |
| --- | --- | --- | --- | --- | --- | --- |
| Aae IAEA (30) | 100% | 96% | 0.24043 | 100% | 88% | 0.0523 |
| Aae LSHTM (30) | 100% | 100% | No Difference | 100% | 80% | **0.0105** |
| Aae UG (15) | 100% | 100% | No Difference | 100% | 75% | **0.0418** |
| Avex Atl (30) | 97% | 85% | 0.1073 | 85% | 75% | 0.3370 |
| Avex Den (30) | 97% | 96% | 0.834 | 82% | 67% | 0.1863 |
| Avex UG (15) | 100% | 83% | 0.1007 | 100% | 61% | **0.0080** |
| Aalbo IAEA (30) | 92% | 75% | 0.0786 | 72% | 39% | **0.0108** |
| Aalbo UG (15) | 100% | 81% | 0.0810 | 81% | 56% | 0.1473 |
| Aalbo Sav (10) | 100% | 80% | 0.1462 | 80% | 40% | **0.0230** |
| Aarab IAEA (16) | 100% | 100% | No difference | 100% | 100% | No difference |
| Aarab ICL (14) | 100% | 100% | No difference | 100% | 92% | 0.2888 |
| Agam G3 | 100% | 93% | 0.1435 | 98% | 78% | **0.0181** |
| LSHTM (30) |  |  |  |  |  |  |
| Agam G3 ICL (20) | 100% | 93% | 0.1435 | 93% | 72% | 0.0844 |
| Transgenic ICL (20) | 100% | 100% | No difference | 100% | 82% | 0.5998 |
| Agam Kis (40) | 100% | 97% | 0.2727 | 88% | 100% | **0.0247** |
| Agam Tia (30) | 100% | 100% | 0.0905 | 100% | 100% | No difference |
| Ngu LSHTM (30) | 100% | 93% | 0.1435 | 93% | 75% | 0.0593 |
| Ngu ICL (20) | 100% | 100% | No difference | 100% | 77% | **0.0244** |
| Keele Ugl (30) | 100% | 100% | No difference | 95% | 87% | 0.2830 |
| As SDA LSHTM (32) | 100% | 100% | No difference | 100% | 100% | No difference |
| As Red LSHTM (30) | 100% | 92% | 0.1169 | 100% | 78% | **0.0069** |
| As Beech LSHTM (30) | 100% | 92% | No difference | 100% | 84% | 0.0235 |
| As STML LSHTM (30) | 100% | 100% | No difference | 100% | 69% | **0.0010** |
| As Dub5 (30) | 100% | 100% | No difference | 100% | 73% | **0.0024** |
| Coq p UG (16) | 100% | 94% | 0.3275 | 100% | 92% | 0.2557 |
| Coq p Sav (14) | 100% | 100% | No difference | 100% | 100% | No difference |
| Coq p Den (15) | 100% | 100% | No difference | 100% | 100% | No difference |
| Coq p Atl (15) | 100% | 100% | No | 100% | 100% | No |

|  |  |  | difference |  |  | difference |
| --- | --- | --- | --- | --- | --- | --- |
| Cq Mu (30) | 100% | 81% | **0.0128** | 71% | 33% | **0.0149** |
| Cq TPRI (20) | 100% | 84% | 0.0655 | 100% | 48% | **0.0002** |
| Cq Sav (12) | 100% | 66% | **0.0300** | 66% | 55% | 0.5895 |
| Cq UG (8) | 100% | 85% | 0.2701 | 75% | 40% | 0.1704 |
| Otaenior Sav (7) | 100% | 60% | 0.0714 | 70% | 35% | 0.2064 |
| Otaenior UG (7) | 100% | 55% | 0.0520 | 55% | 25% | 0.2696 |
| Otriseri Sav (6) | 100% | 60% | 0.0973 | 60% | 0% | 0.0299 |
| Otriseri UG (6) | 100% | 43% | **0.0363** | 43% | 0% | 0.0826 |
| Osollici Sav (7) | 100% | 77% | 0.1937 | 77% | 29% | 0.0830 |
| Osollici UG (6) | 100% | 100% | No difference | 100% | 0% | Not applicable |
| Pferox Sav (6) | 100% | 80% | 0.2689 | 100% | 0% | Not applicable |
| Pferox UG (6) | 100% | 100% | No difference | 100% | 0% | Not applicable |

**Table 4.**
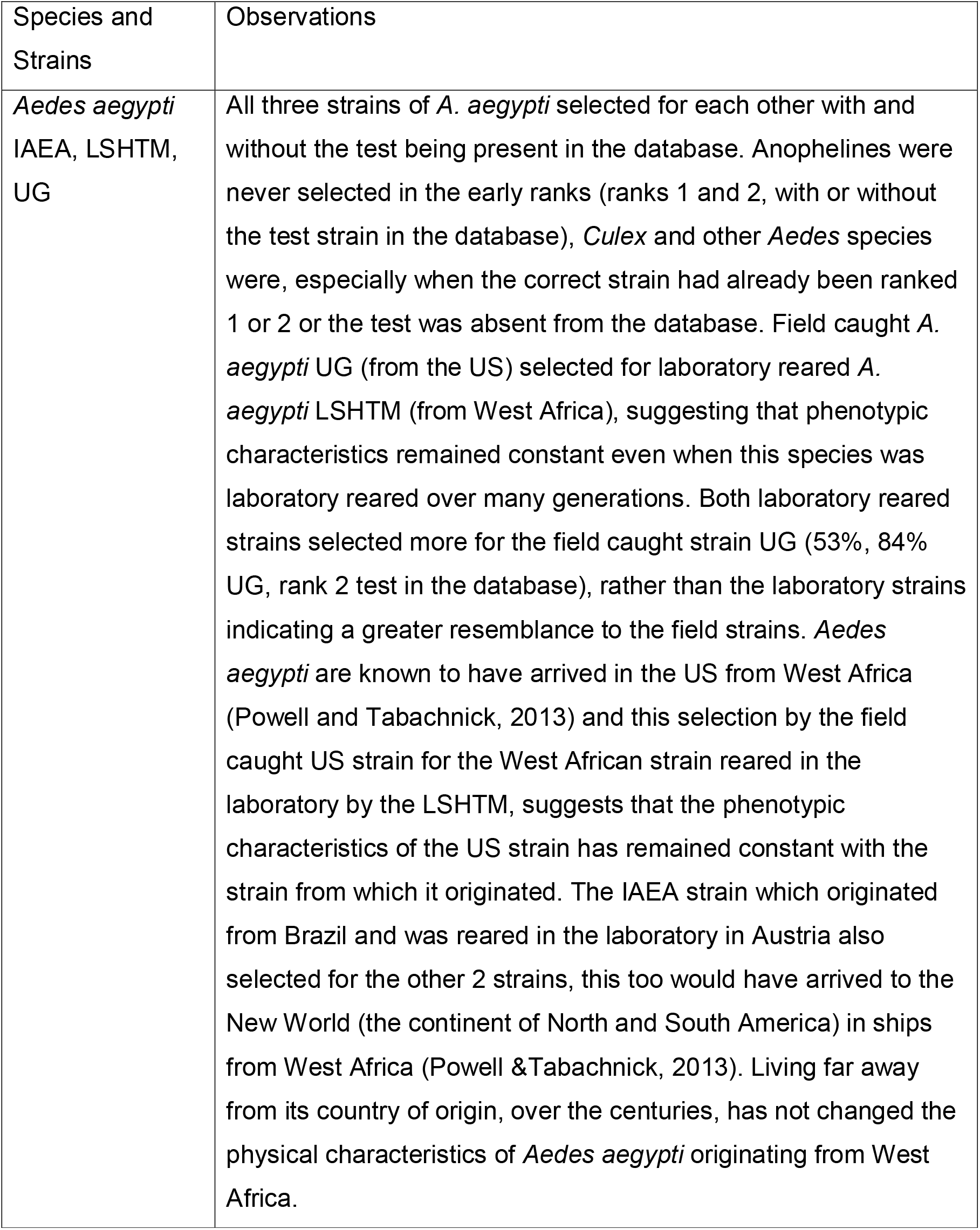
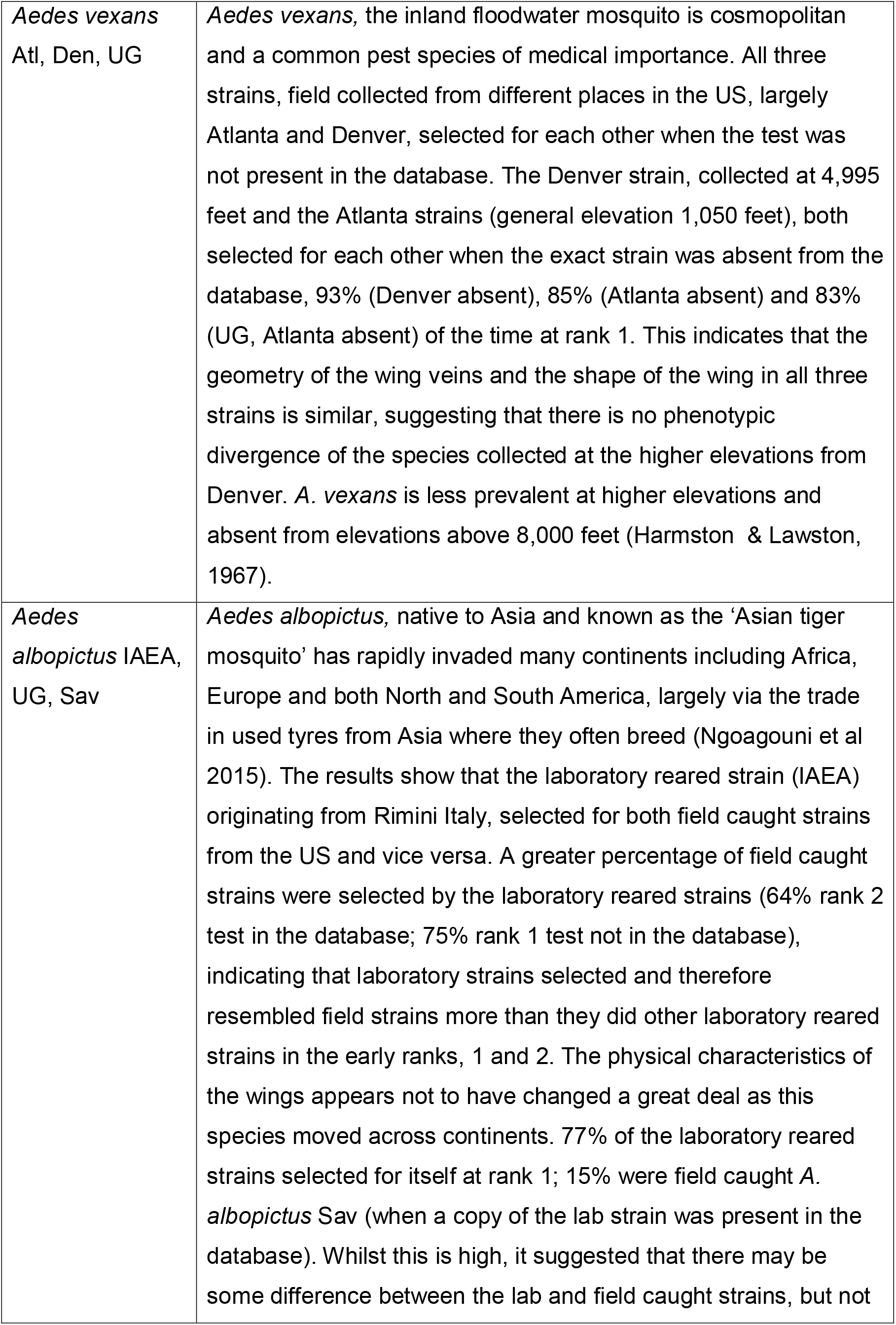
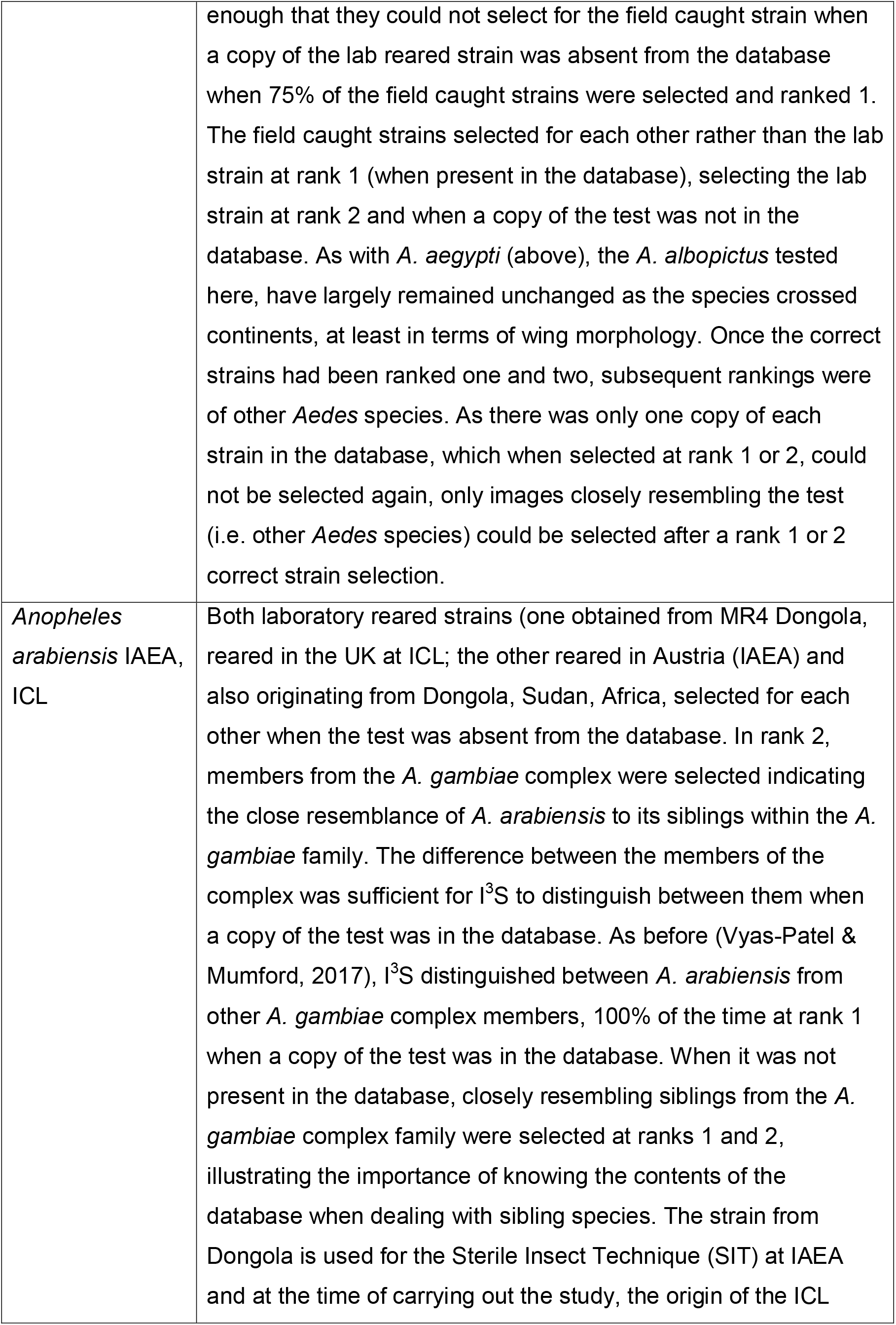
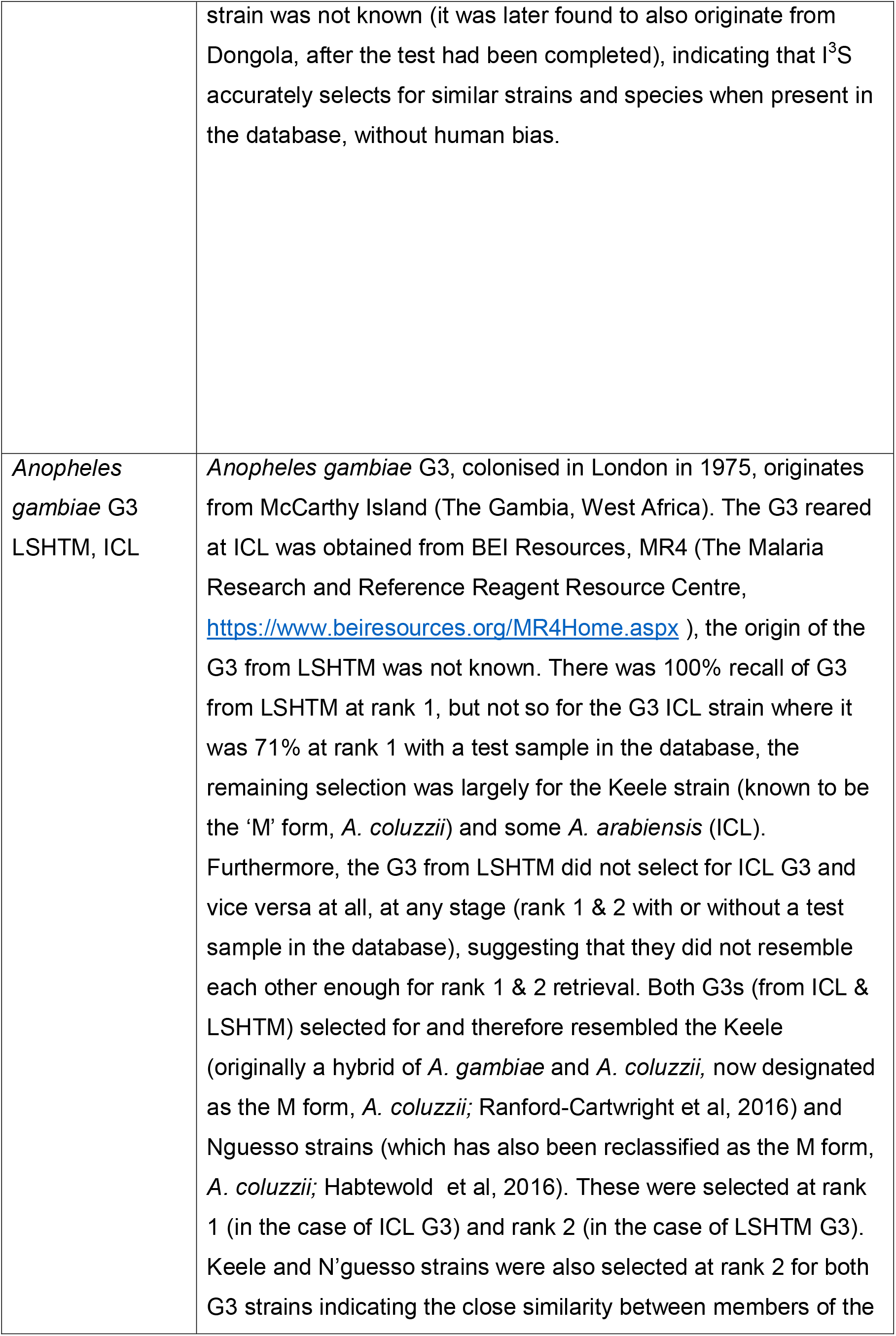
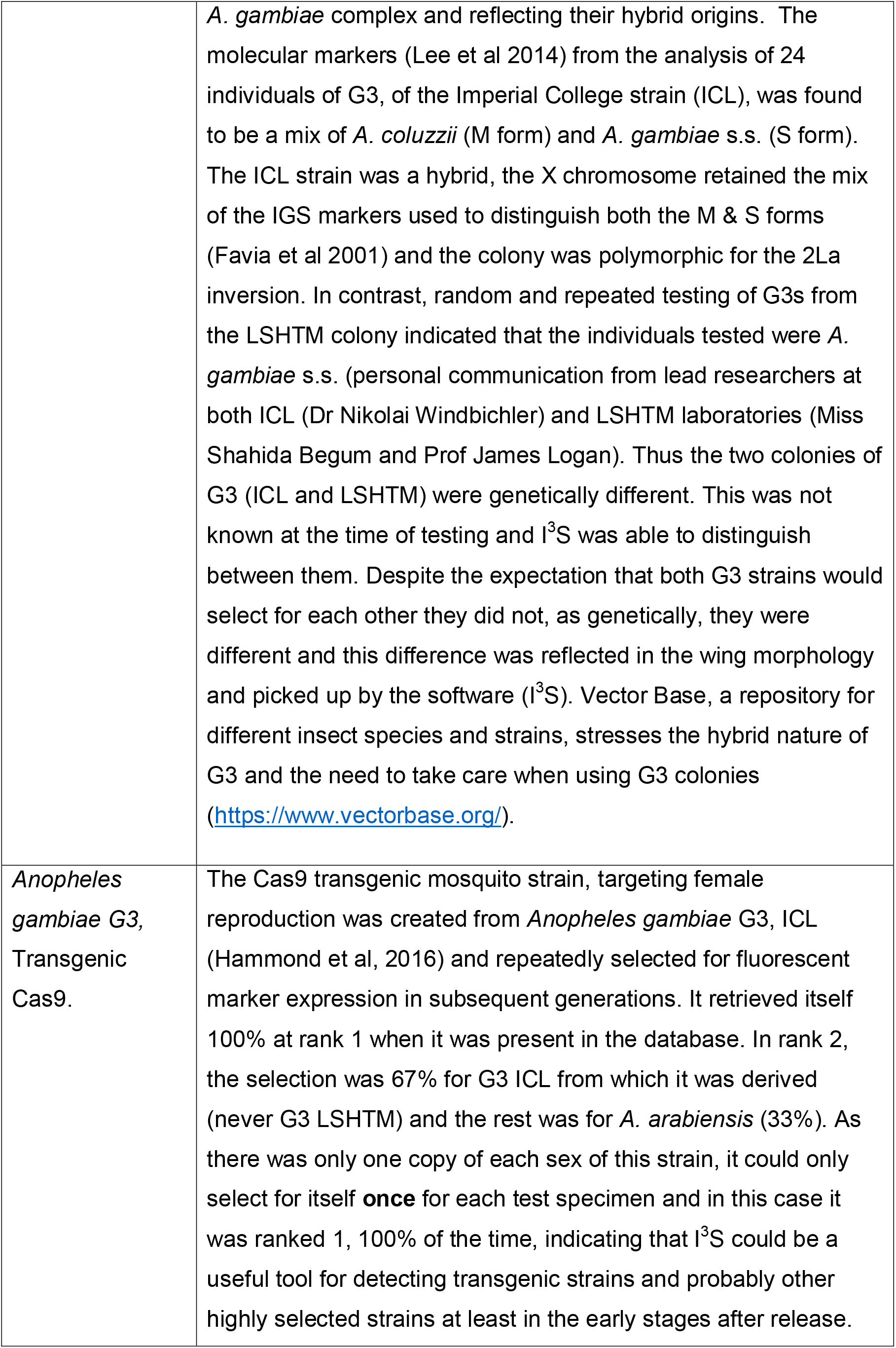
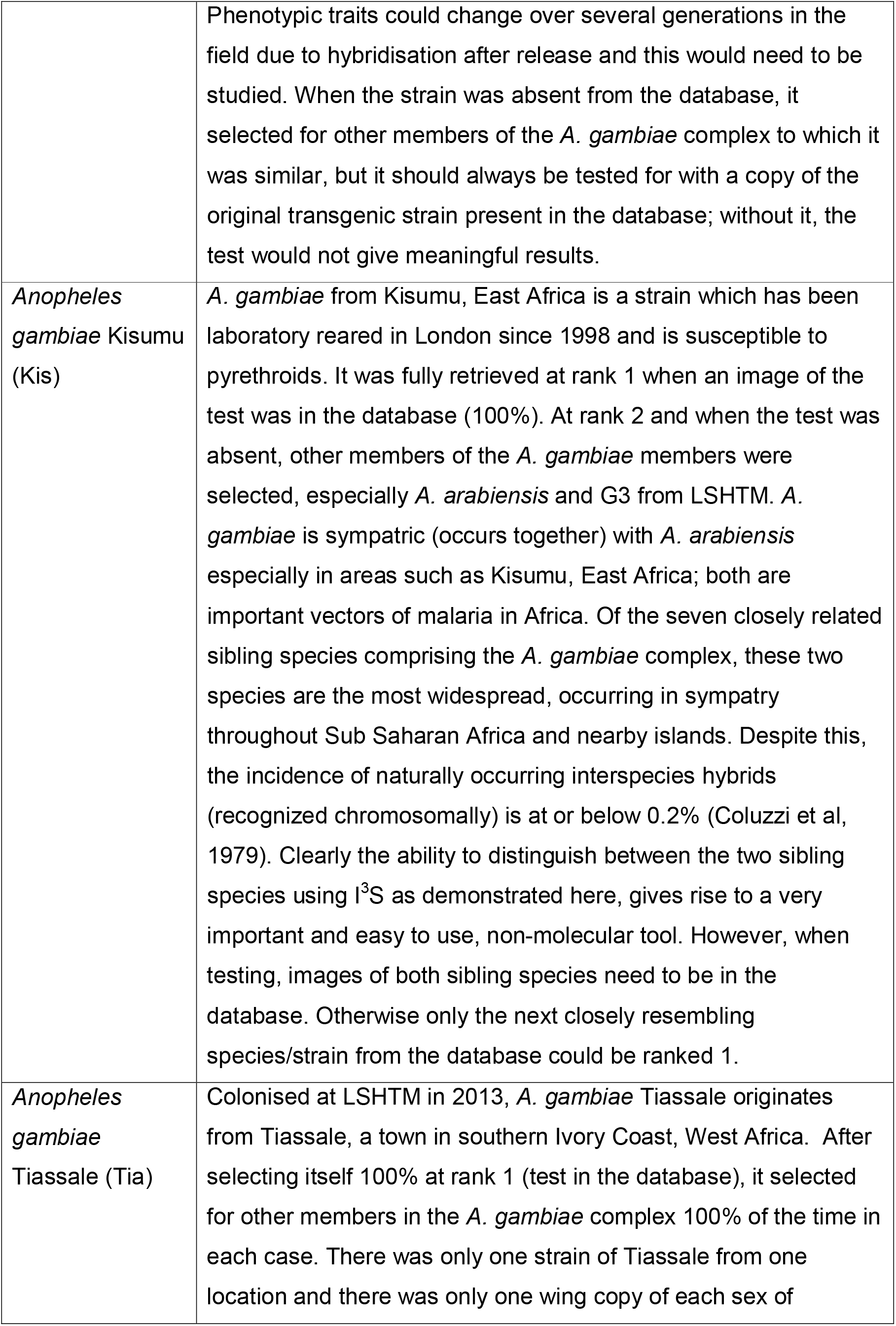
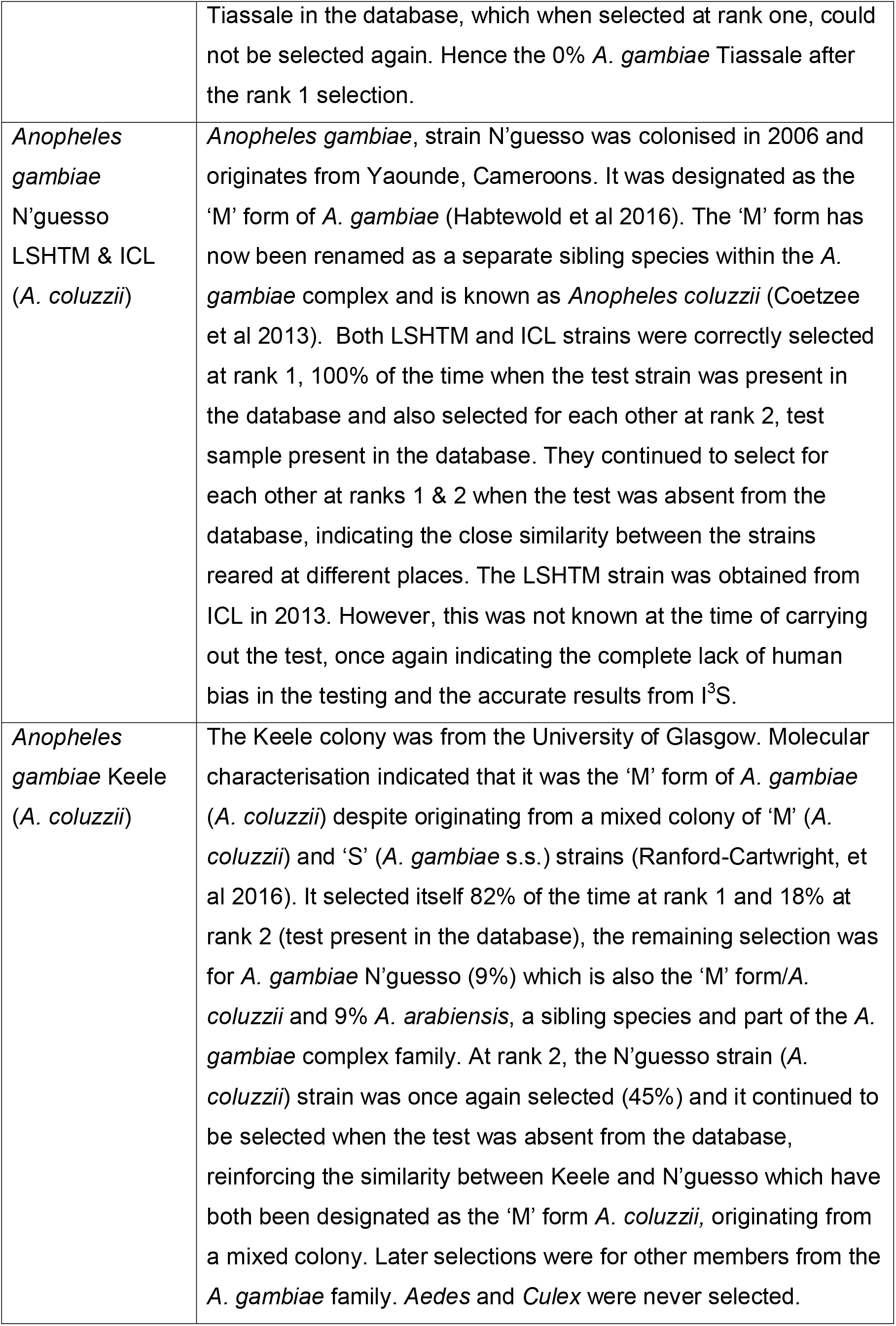
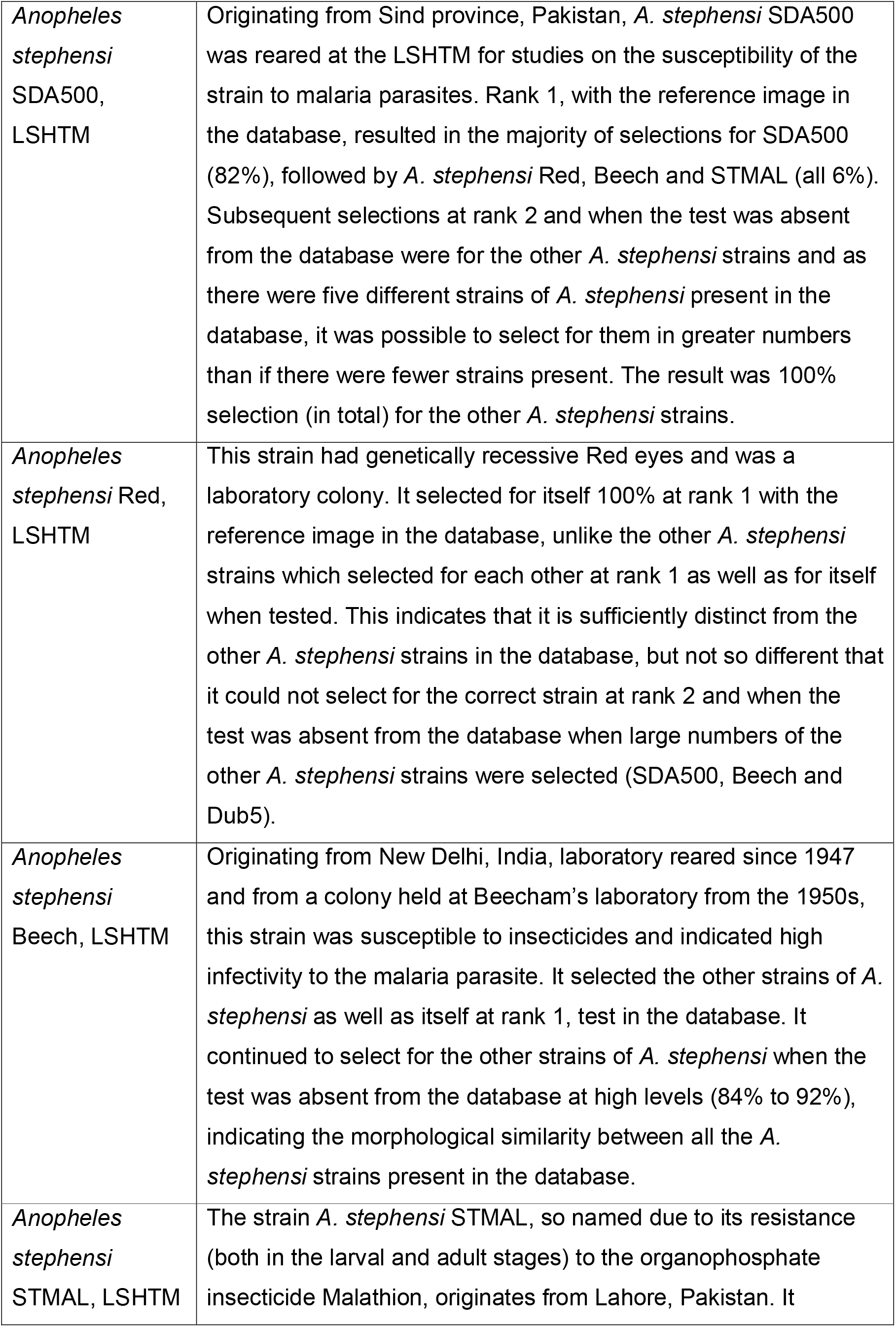
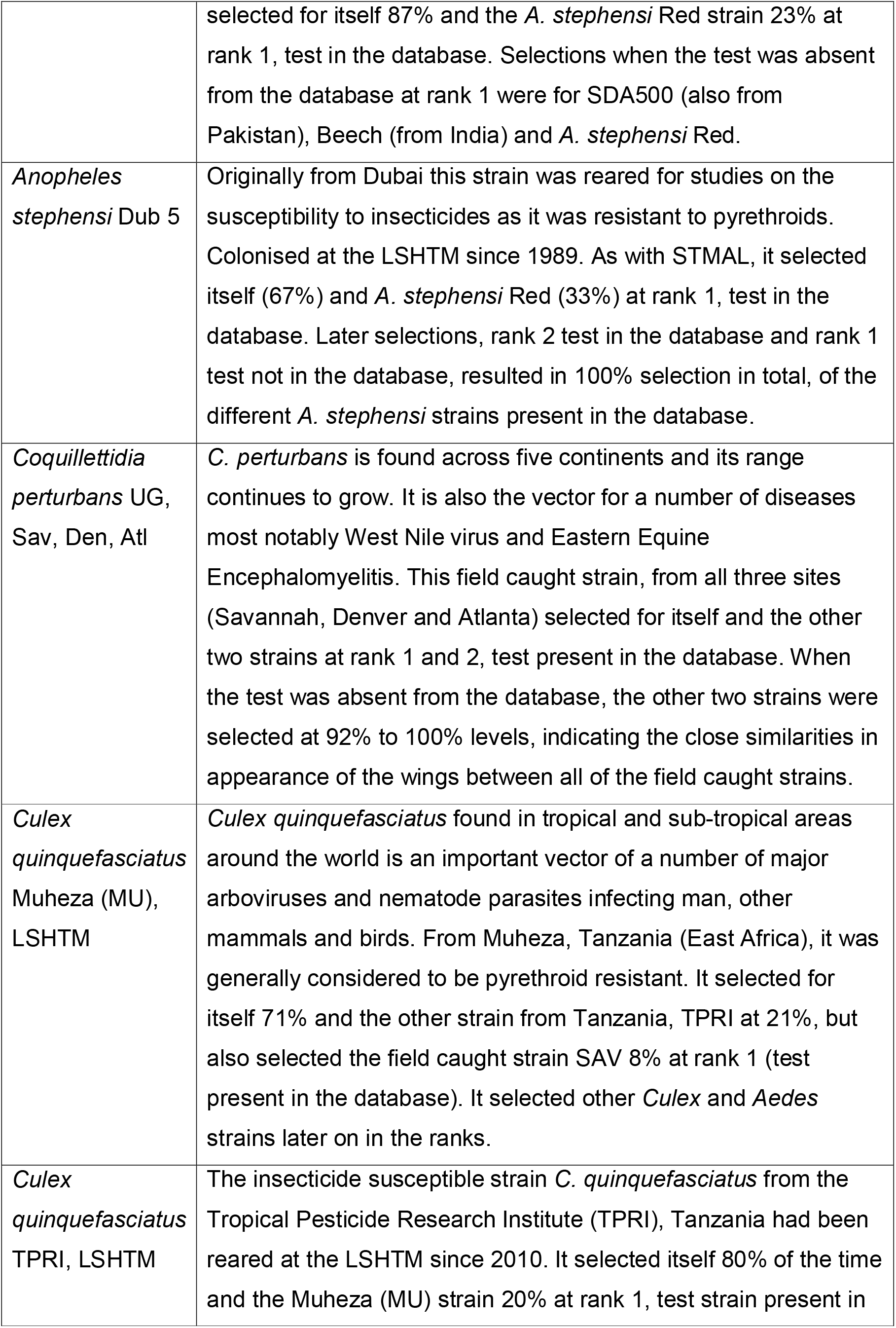
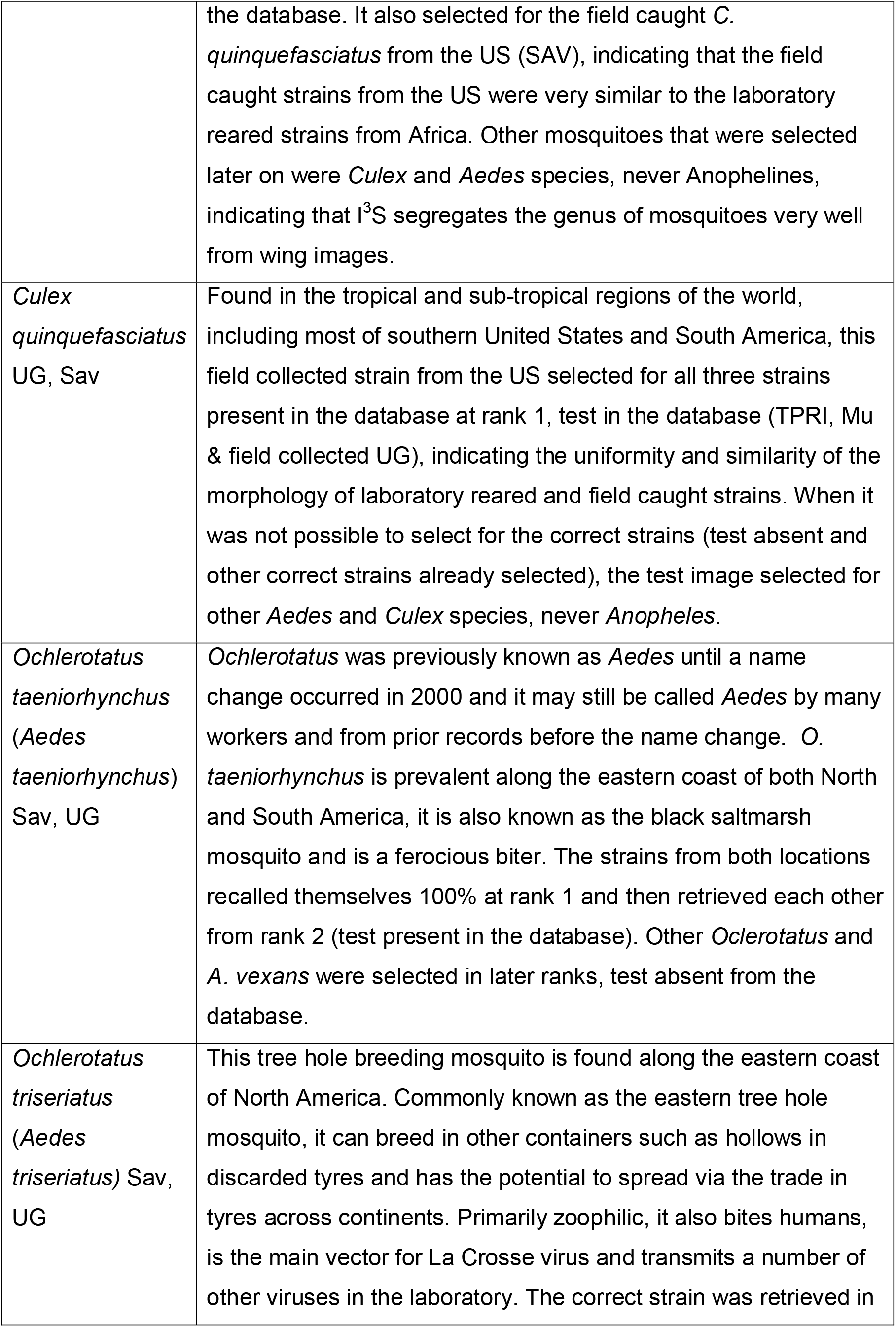
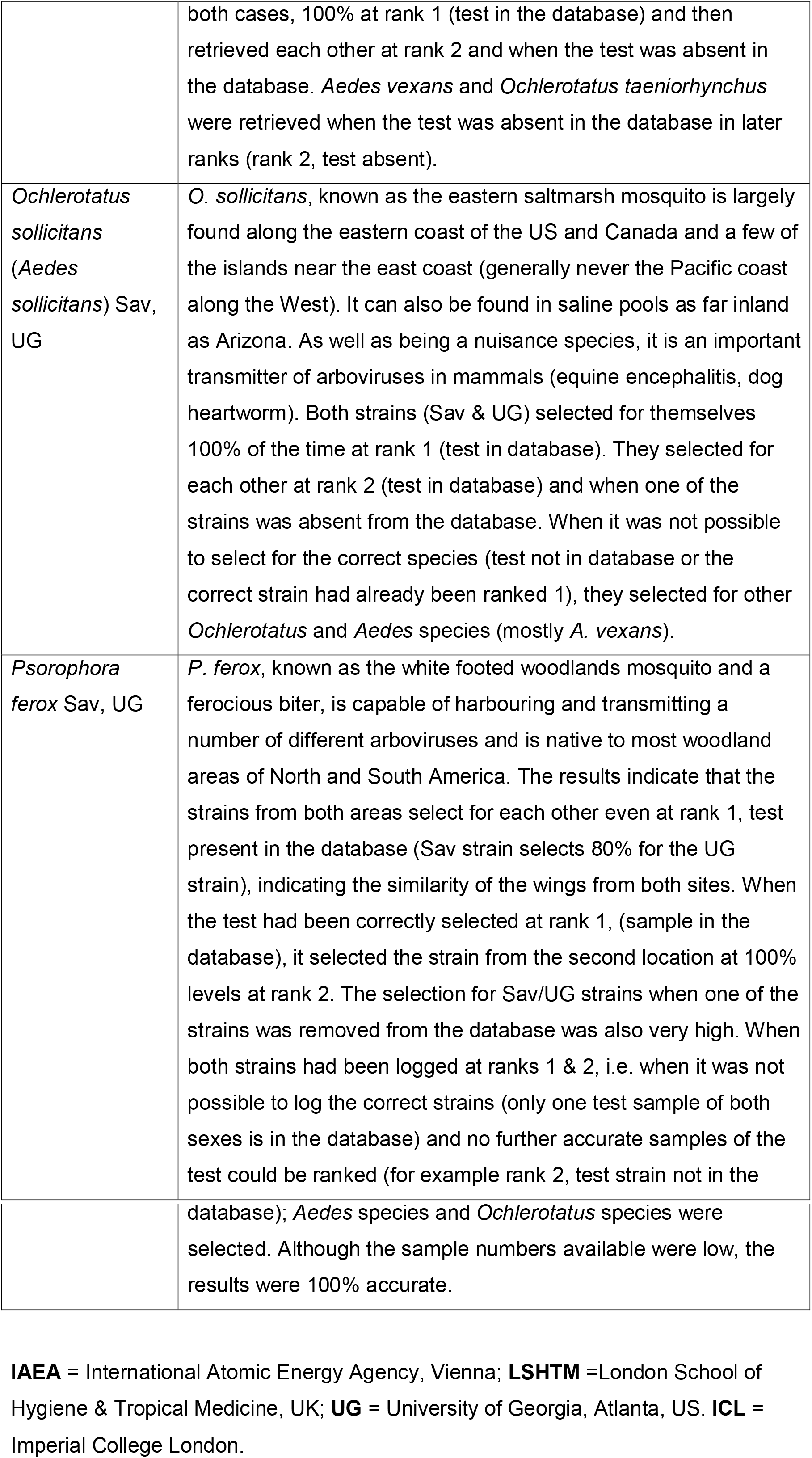
Observations for each species and strain with the test strain either in or out of the database.

## Results

The different mosquito species were identified by the donors, all of them experienced mosquito taxonomists, using traditional mosquito identification keys. Mosquito donors: Centre for Disease Control (CDC) Atlanta (Atl), donor: Dr Rosmarie Kelly; University of Georgia (UG), donor: Dr Elmer Gray; Colorado Mosquito Control CDC, Denver (Den), donor: Dr Michael Weissman; International Atomic Energy Agency IAEA, donor: Dr Jorge Hendrichs; Chatham County Mosquito Control, Savannah (Sav), donor: Dr Laura Peaty & Dr Robert Moulis; Imperial College London, ICL donor: Dr Nikolai Windbichler; London School of Hygiene & Tropical Medicine (LSHTM), donor: Miss Shahida Begum and Prof Mark Rowland. University of Glasgow donor: Dr Lisa Ranford-Cartwright.

The female *Psorophora ciliata* mosquito above, was identified and donated by Dr Elmer Gray, University of Georgia Athens, USA and was field caught in Georgia. The red and blue markings have been exaggerated for clarity. The wings were aligned to be as horizontal as possible with the curved part of the wing to the right hand side and the point of insertion of the wing into the insect thorax to the left; before marking in I^3^S. The blue dots are used by the software for global alignment when comparing two images and were marked first (as dictated by the software). In this study the first blue dot was made at the top of the wing, in line with the kink where the bottom of the wing curves to meet the point of insertion of the wing into the thorax. The second blue dot was made on the far right end of the wing (at the longest point on the curve of the wing). The third is at the bottom of the wing (widest part), so all of them outline the edge of the wing. The red dots were made at points where the veins meet the periphery of the wing and also their junctions within the wing.

The full names of the strains and locations collected are the same as in Tables 1, 2 & 4. The numbers in brackets next to the strains indicate the total number of strains tested. The comparison of proportions was calculated using Med Calc at the 95% confidence interval and 0.05 significance level - Med Calc https://www.medcalc.org/calc/comparison_of_proportions.php. Significant values (p < 0.05) are in bold in the above table.

Not applicable = As there were only 2 similar strains in the database, each strain with just one wing image in the database, when one of the strains was removed it would leave just one wing image, which when selected at rank 1 would leave no further images of the correct strain to be ranked 2 from the database. Hence the 0% result at rank 2 when 100% of the one remaining strain had been ranked at 1.

## Discussion

I^3^S can differentiate between different species of insects using wing images (Vyas-Patel & Mumford, 2017). The present investigation indicated that I^3^S could also retrieve the correct species and strain if it was present in the database with 92% to 100% accuracy at rank 1 and select a different strain of the same species when the test strain was absent from the database (Tables 2, 3, 4). If the exact strain was absent from the database, closely resembling sibling species of the test strain would be selected provided they were present in the database (Table 2, *A. gambiae* species complex). I^3^S is designed so that the image that most closely resembles the test is ranked first (i.e. rank 1), followed by the next most closely resembling image at rank 2 onwards. Concerns that differences in the morphology of strains from different places might affect accurate identifications were therefore unfounded.

Of the forty different strains tested - twenty field caught and twenty laboratory reared, three were significantly different at rank 1 when tested with and without the test specimen in the database (Table 3). In the laboratory reared *C. quinquefasciatus* Muheza (Cq Mu), the reason for the significant result, with a difference of only 19% (Table 3) was not clear. It could simply be different unknown factors that may affect laboratory reared species such as inbreeding, outbreeding (if it was originally a hybrid); or the different conditions, rigours and stress of continuous laboratory rearing in different places. It could be that the Muheza strain was different to the other *C. quinquefasciatus* strains and molecular tests could possibly shed further light on the differences seen here. However, molecular differences were unlikely as the significance level was low and it may simply be the effect of broken wings in one strain affecting the results in other strains. In the field caught *C. quinquefasciatus* Savannah strain (Cq Sav), the numbers were low, with broken wings, this could contribute to the significant difference seen in Table 3, not just for Cq Sav but also for Cq Mu (and the other strains), which could not and did not select for Cq Sav in greater numbers. Similarly in the field caught *O. triseriatus*, where there were also low numbers and imperfect wings, giving rise to the significant result. However low numbers did not automatically mean that accurate results were affected. In the case of field caught *P. ferox* Sav and UG, both low in numbers (6 wings each) but with perfect, unbroken wing samples, the result was 100% selection of each strain for the other when one strain was taken out of the database (Tables 2, 3 and 4).

Of the rank 2 significant results (13 significant out of the 40 strains tested), the higher numbers of significant results were largely due to the fact that the pool of similar strains was twice reduced, firstly because the test strain had been removed from the database and secondly because many of the correct species and strains would already have been selected at rank 1 and therefore could not be selected again as there was only one copy of each strain in the database. Of the 13 significant results at rank 2, 10 were from the laboratory reared samples and only 3 were field caught strains. In this study, field caught strains were similar to each other compared to laboratory reared strains of the same species. Part of the reason for this could be the historical attempts by different workers to mix populations of a species collected from different regions into one cage, in order to prevent inbreeding during laboratory rearing, for example in the case of the Keele strain (Ranford-Cartwright, 2016).

A notable example of the above practice was seen in the results with the laboratory strain of *A. gambiae* G3, reared at both ICL and the LSHTM. The study was carried out with the expectation that the G3 strains from both places would select for each other, but not once did they do so, they were distinct from each other at both ranks 1 and 2. At the time of testing the genotype of the strains tested was not known. Subsequent enquiries revealed that previous molecular characterisation had indicated that the G3 from ICL was a hybrid of the M & S forms of *A. gambiae*, i.e. *A. coluzzii* and *A. gambiae* s.s. (Favia et al 2001) and that from the LSHTM was largely *A. gambiae* s.s. (Table 4). This indicated that the results from I^3^S were fully in keeping with the results of molecular tests. In this study morphological variation followed genetic variation and was picked up by the software, I^3^S.

*A. stephensi*, the other important vector for both *Plasmodium vivax* and *Plasmodium falciparum* malaria parasites, has a wide geographic range from the Middle East to India and continuing into China. Nonetheless, all of the strains tested here selected for each other, with and without the test in the database, with no significant differences at rank 1. Anophelines never selected for Culicines and vice versa indicating the accuracy of I^3^S in selecting the correct genus in the early ranks (ranks 1 & 2).

There was very little strain difference between the samples collected from the field in the US. Any significant difference at rank 1, as occurred with *C. quinquefasciatus* (Sav) and *O. triseriatus* (UG), were probably due to broken wing samples affecting the analysis. The mosquito species collected from the field and tested here do not vary greatly (in terms of wing morphology) from place to place and a species from one area could easily select for the correct species from another area if the test was not in the database. Most notable for its high uniformity in appearance was *Coquillettidia perturbans* which was selected correctly across the board in all cases at 100% or very near 100% levels (92%).

Based on the results seen in this study, it should be possible to use I^3^S to accurately differentiate between hybrid forms of *A. gambiae* in a collection of samples, also the ‘M’ & ‘S’ forms (A. *coluzzii* and *A. gambiae* s.s.) and to separate members of the *A. gambiae* complex with great accuracy. In order to do this, all forms of the *A. gambiae* complex, or at least those forms that were of interest, would need to be in the database, when the correct strain/sibling species was selected 100% of the time. It is no longer the case that ‘’ the M & S forms of *A. gambiae* are morphologically indistinguishable and can only be identified using molecular techniques’’ Lee et al (2014). I^3^S and many other geometric techniques are capable of separating these sibling species and strains and should be used (Cañas-Hoyos et al 2014, Cao et al 2014 and 2015).

I^3^S could also be used to differentiate transgenic or other highly selected mosquitoes from wild ones in the first phase after release, based on the results seen here, where transgenic *A. gambiae* were differentiated and retrieved 100% at rank 1 from the parent species from which they were genetically modified (here it was *A. gambiae* G3, ICL strain). This would require the prior testing of all wings, the transgenic or otherwise selected form, the parent and the wild population wing images and the incorporation of all the relevant wing images into the database. The *A. gambiae* G3 parent tested here is a hybrid strain of *A. gambiae* and the results indicated that it could be distinguished from *A. gambiae* s.s (the LSHTM G3), which occurs in the field. Only one transgenic line was assessed here, so it should .not be assumed that every type of genetically modified mosquito could be similarly discriminated from the parent strain and each new transgenic line should be tested independently. As the parent strain was mostly selected at rank two (67% parent G3 at rank 2 when both transgenic and the parent G3 were in the database), there was a difference between them, albeit a small one. If the difference was large, the majority of the parent strains would have largely been selected beyond rank two, i.e. rank three, four or five. The reasons for any difference in wings between the transgenic and parent mosquitoes may be due to the effects from gene insertions or from selection in the insertion, rearing and maintenance process. Each generation is selected for with fluorescent marker expression, for instance. Furthermore, the exact mechanism and process could be different for any given transgenic mosquito produced, it might be subject to change over a few generations and would need further study. The long term identification potential after transgenic mosquito release is a conjecture, as it is not known what, if any changes could occur in wing morphology after many generations, over the long term, following hybridisation after any transgenic mosquito release.

Several studies (using different methods) have made distinctions between the shape of the wing, the size of the wing and the geometric pattern of the veins on the wing when considering the effects of the environment on morphological change in insect wings (Pieters et al 2017; Liu 2016; Yi Bai et al 2016 and 2015; Demari-Silva 2014; Prudhomme 2016; Sanzana et al 2013; Vidal & Suesdek 2012; Carvajal et al 2016). Here, I^3^S takes into account the shape and vein pattern of the wings, as indicated by the red dots (Figure 1). The software requires the operator to mark the image. This creates a cloud of relevant points. I^3^S only considers the relative positions and distances between these points. In this study, these points represent a set of features such as wing vein branches and wing shape, which are characteristic and present across the range of mosquito species and strains being examined. Apart from the marking of these landmark features on the wing image by the operator, all of the calculations were achieved by the software to arrive at the closest match from the database and the ranking of all the images from the database, according to how closely they matched the test image (information contained on the I^3^S website and associated tutorials http://reijns.com/i3s/, freely downloadable with the software, Hartog & Reijns 2013).

In this study the results relate to mosquito species, but there is every reason to believe that I^3^S could just as effectively be used for other species and strains of insects. Using a variety of different geometric morphometric techniques has resulted in the differentiation of strains in other insect species (Cañas-Hoyos et al 2014, Cao et al 2014 and 2015). A comprehensive review of different geometric methods using mosquitoes as the study insect, described in detail how different geometric morphometric methods could be used to differentiate between insect species and strains, Lorenz et al (2017). The review stressed how all of the geometric morphometric methods reported thus far were capable of sorting not just species but also different strains of mosquitoes and other insects.

Every new species and strain should be tested independently, especially if different methods, software or tools are used. Information about the test species (where it was collected and when) should always be used and recorded. Features other than the wings should also be considered when testing any unknown species against a database of known species.

The results of image recognition software are ‘appearance based’. The software is designed to scrutinise and assess minute details of the markings made of an image, retrieve the closest match for a test from a database and rank the database images according to how closely they resemble the test. The software cannot ascertain the genotype of an organism, however, molecular differences were always reflected in the wings (at least in this study) for example the hybrid and non-hybrid *A. gambiae* G3 strains and could be discriminated and differentiated by the software. I^3^S can sort, differentiate and identify different species and strains of a species with 100% accuracy as long as copies of the test are present in the database. It is a valuable complement to molecular characterisation and is an additional, useful tool to accurately identify species and strains.

One of the strongest reason to continue research into methods that use image recognition and geometric morphometrics techniques, is from a report of the National University of Colombia (Universidad Nacional de Colombia). The group from Colombia, Cañas-Hoyos et al (2014), described how two different strains of *Spodoptera frugiperda*; one that fed on corn and one on rice; could not be differentiated by any type of molecular method, yet they were able to differentiate between the two strains using wing morphometrics and stressed the importance of using the technique in the field. In the cohort of different species/strains examined here, molecular differences (if they existed) between species/strains were always reflected in the wings and could be picked up by the software, note the example of the laboratory reared *A. gambiae* G3 reared at different laboratories one of which was a hybrid and the other *A. gambiae* s.s. Both were differentiated 100% using I^3^S when reference copies were in the database. Based on the examples mentioned above and all of the results seen in this study, it can safely be stated that any molecular difference between different species and strains are expressed in the phenotype of the wings and can be detected using image recognition such as I^3^S or any one of the different geometric morphometric methods available currently.

The case for using image recognition and geometric morphometric tools to identify insect species and strains should not be ignored. These methods should be used alongside molecular characterisation and traditional taxonomic identification using keys. Will and Rubinoff (2004), urged that decisions taken in defining species and strains be based on ‘a constellation of data’ rather than just the narrow findings from any particular discipline, a conclusion supported here. Image recognition software for identifying insect species can clearly complement molecular methods and can be used effectively in conjunction with them or separately. Here, the image recognition software, I^3^S has proven to be an accurate tool in the identification of different species and strains of insects, using their wing images.

## Conclusion

1. A comprehensive study of a large number of different mosquito species and strains indicated that I^3^S was accurately able to detect the differences not just between species but also between different strains of mosquitoes and rank them accordingly.
2. I^3^S could accurately be used to differentiate between members of the *A. gambiae* complex, such as *A. gambiae* s.s., *A. coluzzii* and *A. arabiensis* with 100% accuracy when these sibling species were present in the database.
3. It is no longer the case that molecular characterisation provides the only way to identify sibling species of the *A. gambiae* complex. I^3^S is capable of separating/identifying members of the complex with 100% accuracy.
4. In some cases of strain identification where it was impossible to identify insect strains using molecular characterisation, I^3^S along with other geometric morphometric methods, could be able to differentiate these strains/sibling species and this should always be explored.
5. The presence of hybrid strains could be detected from either laboratory reared or field collected samples, provided a copy of the hybrid in question was present in the database.
6. The transgenic mosquitoes tested in this study could be detected and discriminated from the parent strains, suggesting highly selected lines of mosquitoes can be distinguished from parent strains.
7. I^3^S can accurately be used to identify species and strains of medical importance and is a useful tool to keep watch on potentially invasive species/strains and their spread, such as *Aedes albopictus* from field collections.
8. Field specimens of the same species of mosquitoes collected from geographically different areas from the US vary little from each other and I^3^S is capable of accurately selecting other strains of the test species if the test strain is absent from the database.
9. Laboratory reared strains can vary from different laboratories (as indicated by I^3^S for example *A. gambiae* G3). Both morphological and molecular tests to identify the exact strain would be prudent before their use in experimental studies.
10. I^3^S makes for an excellent additional and accurate tool to be used with both molecular and traditional identification using keys and should be utilised.
11. In future keeping copies of insect wings and other insect body parts, in databases, for use in image recognition of species/strain identification could become the norm.

## Acknowledgements

Without the generous donation of insects from everyone mentioned in Table 1 (mosquito donors) this work would not have been possible. The interest and encouragement, correspondence and good wishes from every mosquito donor, was vital for the completion of this project. The correspondence with the I^3^S team, Jurgen Hartog and Renate Reijns, was similarly invaluable. Thank you all!

